# Artificial Intelligence-rationalized balanced PPARα/γ dual agonism resets the dysregulated macrophage processes in inflammatory bowel disease

**DOI:** 10.1101/2021.07.18.452807

**Authors:** Gajanan D. Katkar, Ibrahim M. Sayed, Mahitha Shree Anandachar, Vanessa Castillo, Eleadah Vidales, Daniel Toobian, Fatima Usmani, Joseph R. Sawires, Geoffray Leriche, Jerry Yang, William J. Sandborn, Soumita Das, Debashis Sahoo, Pradipta Ghosh

**Author notes:** **Alternative affiliation:** Department of Medical Microbiology and Immunology, Faculty of Medicine, Assiut University, Egypt (Ibrahim M. Sayed). **Correspondence to**: **William J. Sandborn, M.D.;** Professor, Department of Medicine, University of California San Diego; 9500 Gilman Drive, MC 0956, La Jolla, CA 92093-0831., **Soumita Das, Ph.D.;** Associate Professor, Department of Pathology, University of California, San Diego; 9500 Gilman Drive, George E. Palade Bldg, Rm 256, 239; La Jolla, CA 92093. : **Email:**, **Debashis Sahoo, Ph.D;** Assistant Professor, Department of Pediatrics, University of California San Diego; 9500 Gilman Drive, MC 0730, Leichtag Building 132; La Jolla, CA 92093-0831., **Pradipta Ghosh, M.D.;** Professor, Departments of Medicine, and Cell and Molecular Medicine, University of California San Diego; 9500 Gilman Drive (MC 0651), George E. Palade Bldg, Rm 232, 239; La Jolla, CA 92093.

## Abstract

A computational platform, the Boolean network explorer (*BoNE*), has recently been developed to infuse AI-enhanced precision into drug discovery; it enables querying and navigating invariant Boolean Implication Networks of disease maps for prioritizing high-value targets. Here we used *BoNE* to query an Inflammatory Bowel Disease (IBD)-map and prioritize a therapeutic strategy that involves dual agonism of two nuclear receptors, PPARα/γ. Balanced agonism of PPARα/γ was predicted to modulate macrophage processes, ameliorate colitis in network-prioritized animal models, ‘reset’ the gene expression network from disease to health, and achieve a favorable therapeutic index that tracked other FDA-approved targets. Predictions were validated using a balanced and potent PPARα/γ-dual agonist (PAR5359) in two pre-clinical murine models, i.e., *Citrobacter rodentium*-induced infectious colitis and DSS-induced colitis. Using a combination of selective inhibitors and agonists, we show that balanced dual agonism promotes bacterial clearance more efficiently than individual agonists, both *in vivo* and *in vitro*. PPARa is required and its agonism is sufficient to induce the pro-inflammatory cytokines and cellular ROS, which are essential for bacterial clearance and immunity, whereas PPARg-agonism blunts these responses, delays microbial clearance and induces the anti-inflammatory cytokine, IL10; balanced dual agonism achieved controlled inflammation while protecting the gut barrier and ‘reversal’ of the transcriptomic network. Furthermore, dual agonism reversed the defective bacterial clearance observed in PBMCs derived from IBD patients. These findings not only deliver a macrophage modulator for use as barrier-protective therapy in IBD, but also highlight the potential of *BoNE* to rationalize combination therapy.

## INTRODUCTION

Inflammatory bowel disease (IBD) is an autoimmune disorder of the gut in which diverse components including microbes, genetics, environment and immune cells interact in elusive ways to culminate in overt diseases (1–3). It is also heterogeneous with complex sub-disease phenotypes (i.e., strictures, fistula, abscesses and colitis-associated cancers) (4, 5). Currently, patients are offered anti-inflammatory agents that have a ~30-40% response-rate, and 40% of responders become refractory to treatment within one year (6, 7). Little is known to fundamentally tackle the most widely recognized indicator/predictor of disease relapse i.e., a compromised mucosal barrier. Homeostasis within this mucosal barrier is maintained by our innate immune system, and either too little or too much reactivity to invasive commensal or pathogenic bacteria, is associated with IBD(8). Although defects in the resolution of intestinal inflammation have been attributed to altered monocyte–macrophage processes in IBD, macrophage modulators are yet to emerge as treatment modalities in IBD (8).

We recently developed and validated an AI-guided drug discovery pipeline that uses large transcriptomic datasets (of the human colon) to build a Boolean network of gene clusters (9) (**Figure 1**; *Step 0*); this network differs from other computational methods (e.g., Bayesian and Differential Expression Analyses) because gene clusters here are interconnected by directed edges that represent Boolean Implication Relationships that invariably hold true in every dataset within the cohort. Once built, the network is queried using machine learning approaches to identify in an unbiased manner which clusters most effectively distinguish healthy from diseased samples and do so reproducibly across multiple other cohorts (906 human samples, 234 mouse samples). Gene-clusters that maintain the integrity of the mucosal barrier emerged as the genes that are invariably downregulated in IBD, whose pharmacologic augmentation/induction was predicted to ‘reset’ the network. These insights were exploited to prioritize one target, choose appropriate pre-clinical murine models for target validation and design patient-derived organoid models (**Figure 1**; *Step 0*) (9). Treatment efficacy was confirmed in patient-derived organoids using multivariate analyses. This AI-assisted approach provided a first-in-class epithelial barrier-protective agent in IBD and predicted Phase-III success with higher accuracy over traditional approaches (9).

**Figure 1.**
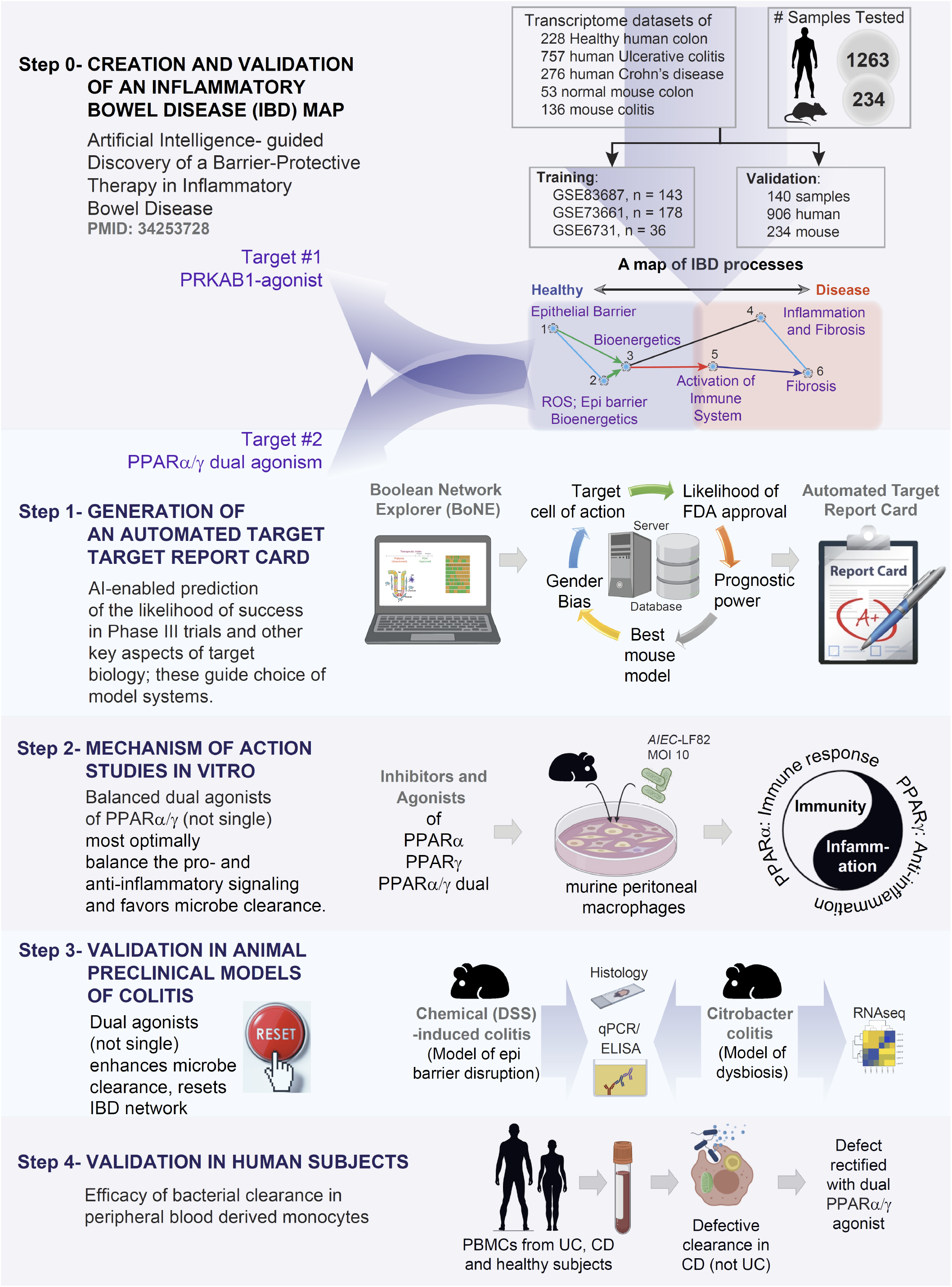
Study design. *From top to bottom:* The premise of a 4-step drug discovery pipeline is summarized on the top (Step 0) is a recently published(9) Boolean implication network-based computational model of disease continuum states in inflammatory bowel disease (IBD map). The map, comprised of 6 gene clusters, was created and validated database containing 1497 gene–expression data (1263 human and 234 mouse samples). Paths, clusters and a list of genes in the network-based model were prioritized to discover one clinically actionable drug target (*PRKAB1*)(9). Steps 1-4 outline the AI-guided identification and validation of another target pair, *PPARA* and *PPARG*. Step 1: Dual agonists of PPARα/γ were predicted to—(i) modulate epithelial and macrophage processes; (ii) *Citr<u>o</u>bacter* and chemical models of colitis were predicted as most optimal models; (iii) have high therapeutic index indicative of likelihood to succeed in Phase III clinical trials. Step 2: A combination of inhibitor and agonist studies helped establish that dual agonists reduce inflammation (PPARg) while ensuring the induction of adequate immune response (PPARα). Step 3: Dural agonists ameliorated colitis in two preclinical models of colitis, and reversed the patterns of disease-associated gene expression that were altered in the IBD map. Step 4: In phase ‘0’ human pre-clinical trials, PBMCs from CD, but not UC or healthy showed defective microbe clearance; this defect was reversed with a dual agonist of PPARα/γ.

Here we use the same AI-guided drug discovery pipeline to identify and validate a first-in-class macrophage modulator that is predicted to restore mucosal barrier and homeostasis in IBD (**Figure 1**; *Steps 1*). Using primary peritoneal macrophages and specific agonists and antagonists, we reveal the mechanism(s) of action that enable balanced agonists of this pair of nuclear receptors, PPARα/γ, to reverse some of the fundamental imbalances of the innate immune system in IBD, such that immunity can be achieved without overzealous inflammation (**Figure 1**; *Step 2*). We demonstrate the accuracy and predictive power of this network-rationalized approach and reveal the efficacy of balanced dual agonists of PPARα/γ in two pre-clinical murine models (**Figure**; *Step 3*) and in patient-derived PBMCs (**Figure**; *Step 4*).

## RESULTS

### Development of a web-based platform for generating a ‘target report card’

We first developed an interactive, user-friendly web-based platform that allows the querying of our Boolean network-based-IBD map with the goal of enabling researchers to pick high-value targets (9) (**Figure 1**; *Step 0;* **Supplemental Figure 1**). The platform generates a comprehensive automated report containing actionable information for target validation, a ‘target report card’, which contains predictions on five components (**Figure 2A**): (i) *Impact on the outcome of IBD in response to treatment*, which shows how levels of expression of any proposed target gene(s) relates to the likelihood of response to therapies across diverse cohorts; (ii) *Therapeutic index*, a computationally generated index using Boolean implication statistics which provides a likelihood score of indicate whether pharmacologic manipulation of the target gene(s) would lead to success in Phase III clinical trials; (iii) *Appropriateness of preclinical mouse models*, a component that indicates which murine models of colitis shows the most significant change in the target genes (and hence, likely to be best models to test the efficacy of any manipulation of that target); (iv) *Gender bias*, a component that indicates whether the gene is differentially expressed in IBD-afflicted men *versus* women; and (v) *Target tissue/cell type specificity*, which shows the likely cell type where the target is maximally expressed, and hence, the cell type of desirable pharmacologic action. Details of how therapeutic index is computed are outlined in *Methods* and in **Supplemental Figure 2**; it is essentially a statistical score of how tightly any proposed target gene(s) associates with FDA-approved targets *versus* those that failed, and serves as an indicator of likelihood of success (9). Similarly, details of how cell type of action is computer are outlined in *Methods* and in **Supplemental Figure 3**.

**Figure 2:**
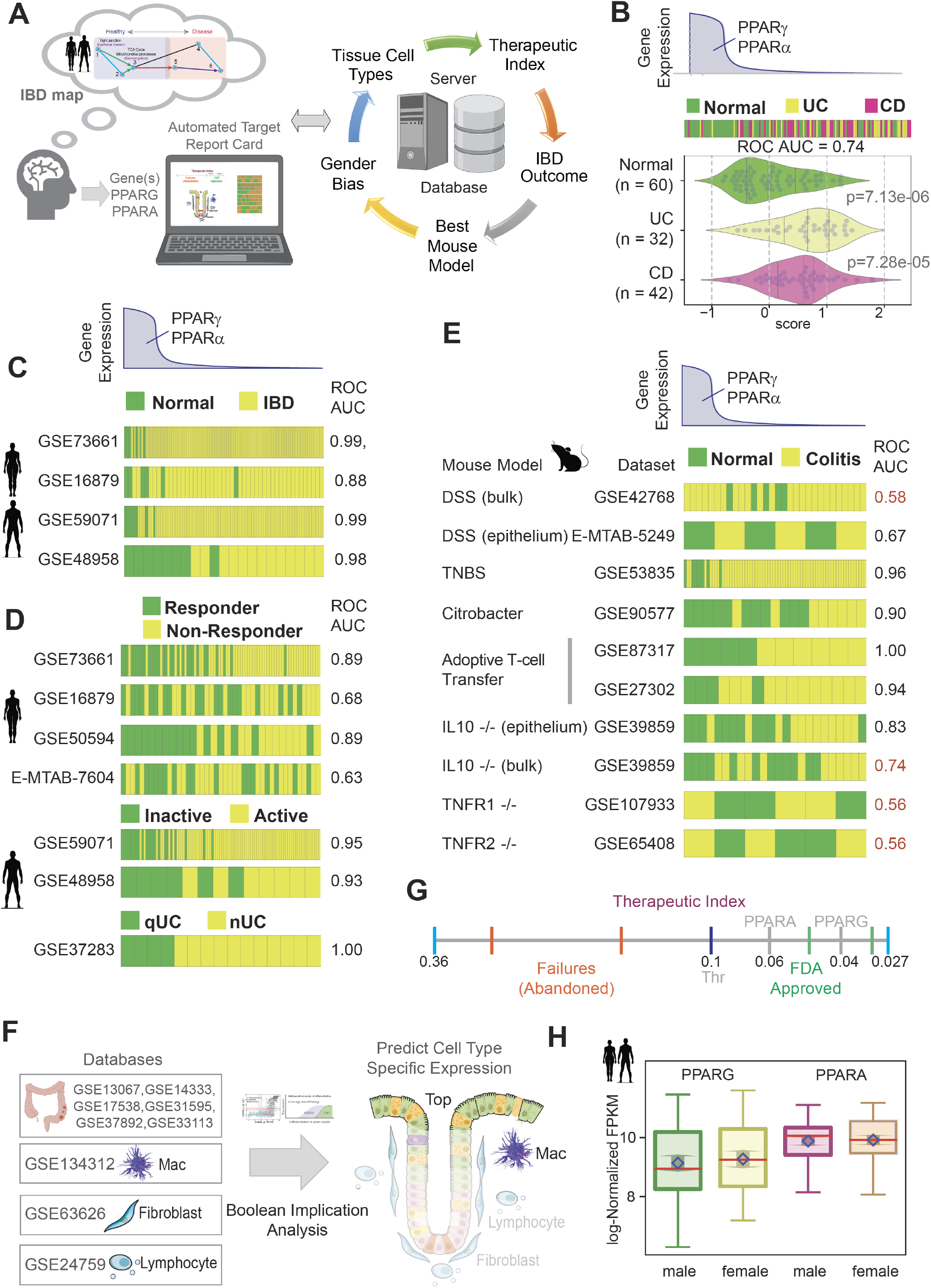
Network-guided rationalization of PPARA/PPARG as targets in IBD. (**A**) An interactive web-based platform allows the querying of paths of gene clusters in the IBD map [(9); see **Supplementary Fig. 1**] to pick high-value targets with a few mouse clicks and generate a comprehensive automated target ‘report card’. The components of a ‘target report card’ is shown (*right*): predicted ‘therapeutic index’ (likelihood of Phase III success), IBD outcome (prognostic potential in UC and/or CD), network-prioritized mouse model, estimation of gender bias and predicted tissue cell type of action. (**B-H**) Components of a target report card for *PPARA* and *PPARG* are displayed. Bar plot (B; top) displays the rank ordering of normal *vs*. ulcerative colitis (UC) /Crohn’s Disease (CD) patient samples using the average gene expression patterns of the two genes: *PPARG/PPARA*. Samples are arranged from highest (left) to lowest (right) levels. ROC-AUC statistics were measured for determining the classification strength of normal vs IBD. Bar plots (B; top) and violin plots (B; bottom) display the differences in the average expression of the two genes in normal, UC and CD samples in the test cohort that was used to build the IBD-map in (9). Bar plots in panel C-D show the rank ordering of either normal *vs*. IBD samples (C) or responder *vs*. non-responder (R *vs*. NR; D), or active *vs*. inactive disease, or neoplastic progression in quiescent UC (qUC *vs*. nUC; D) across numerous cohorts based on gene expression patterns of *PPARG* and *PPARA*, from highest (left) to lowest (right) levels. Classification strength within each cohort is measured using ROC-AUC analyses. Bar plots in panel E show the rank ordering of either normal vs IBD samples across numerous published murine models of IBD based on gene expression patterns of *PPARG* and *PPARA* as in D. ACT = adoptive T cell transfer. Classification strength within each cohort is measured using ROC-AUC analyses. Bulk = whole distal colon; epithelium = sorted epithelial cells. Schematic in F summarizes the computational prediction of the cell type of action for potential PPARA/G targeted therapy, as determined using Boolean implication analysis. GSEID# of multiple publicly available databases of the different cell types and colorectal datasets used to make sure predictions are cited. Red boxes/circles denote that *PPARA/G*-targeted therapeutics are predicted to work on monocytes/macrophages and crypt-top enterocytes. Computationally generated therapeutic index (see *Methods*) is represented as a line graph in G. The annotated numbers represent Boolean implication statistics. *PPARA* and *PPARG* align with other targets of FDA approved drugs on the right of threshold (0.1). Two FDA approved targets (green; *ITGB1*, 0.046; *JAK2*, 0.032), two abandoned targets (red; *SMAD7*, 0.33; IL11, 0.16), *PPARA* (grey, 0.064), *PPARG* (grey, 0.04), and the threshold (black, 0.1) are shown in the scale. Box plot in panel H shows that the level of *PPARA/G* expression is similar in the colons of both genders in health and in IBD, and hence, PPARα/γ-targeted therapeutics are predicted to have little/no gender predilection.

### Ppar-α/γ dual agonists are predicted to be effective barrier-protective agents in IBD

Previous work had identified a little over 900 genes in 3 clusters (Clusters #1-2-3 within the IBD map; **Figure 1**: *Step 0;* **Supplemental Figure 1A-B**) as potentially high-value targets, all of which were invariably downregulated in IBD-afflicted colons (9). Reactome analyses showed that epithelial tight junctions (TJs), bioenergetics, and nuclear receptor pathway (PPAR signaling) related genes that are responsible for colon homeostasis are the major cellular processes regulated by these genes (**Supplemental Figure 1B**). Downregulation of genes in clusters #1-3 was invariably associated also with an upregulation of genes in clusters #4-5-6; reactome analyses of the latter showed cellular processes that concern immune cell activation, inflammation and fibrosis, which are hallmarks of IBD (**Supplemental Figure 1B**). Of the druggable candidates within C#1-2-3, 17 targets were identified as associated with GO biological function of ‘response to stress’/’response to stimuli’. Targeting one of the 17 targets, *PRKAB1*, the subunit of the heterotrimeric AMP-kinase engaged in cellular bioenergetics and stress response successfully restored the gut barrier function and also protected it from collapse in response to microbial challenge (9). Here, we prioritized two more of those 17 targets, *PPARA* and *PPARG*, which encode a pair of nuclear receptors, Ppar-α and Ppar-γ, respectively. These two stress/stimuli-responsive genes are equivalent to each other and to *PRKAB1*, and like *PRKAB1*, are invariably downregulated in all IBD samples (**Supplemental Figure 1B-D**). *PPARA* is in cluster #2 and *PPARG* is in cluster #3 (**Supplemental Figure 1B**). They both were located on the two major Boolean paths associated with epithelial barrier and inflammation/fibrosis (**Supplemental Figure 1B**)(9). Together, these findings imply three things: (i) that *PPARA* and *PPARG* are simultaneously downregulated in IBD, (ii) that such downregulation is invariably associated with inflammation, fibrosis and disruption of the epithelial barrier, and (iii) that simultaneous upregulation of *PPARA* and *PPARG* with agonists may restore the gut barrier. The last point is particularly important because PPARα/γ agonists are known to augment the expression of *PPARA* and *PPARG*, and depletion of either reduced the expression of the other (10).

Noteworthy, while the role of Ppar-g in colitis has been investigated through numerous studies over the past 3 decades (11–13) (**Supplemental Table 1**), the role of Ppar-α has been contradictory (**Supplemental Table 2**), and their dual agonism in IBD has never been explored. All studies agree that Ppar-g agonists ameliorate DSS-induced colitis (13–15). Although claimed to be effective on diverse cell types in the gut (epithelium, T-cells, and macrophages), the most notable target cells of Ppar-g agonists are macrophages and dendritic cells (16). Furthermore, Phase I and II clinical trials with Ppar-g agonists either alone (17, 18) or in combination with mesalamine (19) show barrier protective effects in UC patients. Despite these insights, the biopharmaceutical industry has not been able to harness the beneficial impact of this major target within emergent therapeutic strategies largely due to a trail of withdrawals after devastating long-term side effects including heart failure, bone fracture, bladder cancer, fluid retention and weight gain (20, 21). Intriguingly, and of relevance to this work, the addition of Ppar-α agonistic activity to Ppar-g, Ppar-g, to Ppar-δ agonists have led to a higher safety profile, leading to their development for use in many diseases, including type 2 diabetes, dyslipidemia and non-alcoholic fatty liver disease (22).

### An automated target ‘report card’ for *PPARA* and *PPARG* in IBD

We next generated an automated target report card for *PPARA* and *PPARG*. A high level of both PPARs, determined using a composite score for the abundance of both transcripts, was sufficient to distinguish healthy from IBD samples, not just in the test cohort that was used to build the IBD-map (ROC AUC of 0.74; **Figure 2B;** see also **Supplemental Figure 2A-D**), but also in four other independent cohorts with ROC AUC consistently above 0.88 (**Figure 2C**). High levels of both PPARs also separated responders from non-responders receiving TNFα-neutralizing mAbs, GSE16879, E-MTAB-7604 or Vedolizumab that block the α4β7 integrin to prevent selective gut inflammatory, GSE73661 (ROC AUC 0.63-0.89, **Figure 2D**), inactive disease from active disease (two independent cohorts ROC AUC above 0.93; **Figure 2D**), and quiescent UC that progressed, or not to neoplasia (ROC AUC=1.00 for qUC *vs*. nUC; **Figure 2D**). High level of *PPARA* and *PPARG* was also able to distinguish healthy from diseased samples in diverse murine models of colitis (**Figure 2E**); but such separation was most effectively noted in some models (*Citrobacter* infection-induced colitis, adoptive T-cell transfer, TNBS and *IL10^-/-^*), but not in others (DSS, and *TNFR1/2^-/-^*). These findings imply that therapeutics targeting these two genes are best evaluated in the murine models that show the most consistent decrease in the gene expression, e.g., *Citrobacter* infection-induced colitis, adoptive T-cell transfer, TNBS, etc. This was intriguing because the majority (~90%) of the published work on PPARa/g dual agonists have been carried out in DSS models (**Supplemental Table 1-2**).

The expression profile of the target genes in the gut mucosa revealed that *PPARA* and *PPARG* are co-expressed at the highest levels in the crypt top epithelial cells and macrophages (**Figure 2F; Supplemental Figure 3**), predicting that dual agonists are likely to preferentially act on these two cell types. The therapeutic index was below 0.1 for both genes (0.06 for *PPARA* and 0.04 for *PPARG;* **Figure 2G; Supplemental Figure 2E-F**), aligned well with two other FDA-approved targets shown on the line graph (ITGB1, 0.046 and JAK2, 0.032). The index, which is a statistical measure of the strength of association of Pparα/γ with genes that are targets of FDA-approved drugs that have successfully moved through the three phases of drug discovery (i.e., proven efficacy, with acceptable toxicity). A low number is indicative of a high likelihood of success in Phase-III trials. Finally, *PPARA* and *PPARG* expression was downregulated to a similar extent in men and women with IBD (**Figure 2H**), predicting that therapeutics targeting them are likely to be effective in both genders.

### Rationalization of *PPARA/G* and *PPARGC1A* as targets in IBD

Because proteins, but not transcripts, are the targets of therapeutic agents, the impact of therapeutics is translated to cellular processes *via* protein-protein interaction (PPI) networks, a.k.a interactomes. We next asked how dual agonists of Ppar-α/γ might impact cellular pathways and processes. A PPI network visualized using Ppar-α and Ppar-γ as ‘query/input’ and the interactive STRING v11.0 database (https://string-db.org/) as a web resource of known and predicted protein–protein interactions curated from numerous sources, including experimental data, computational prediction methods and public text collections. Pgc1a (a product of the gene *PPARGC1A*) was a common interactor between the two PPARs (**Figure 3A**). We noted that Pgc1a also happens to be a major component within the Ppar-α/γ functional network, serving as a central hub for positive feedback loops between the PPARs and their biological function (**Figure 3B**), i.e., mitochondrial biogenesis, DNA replication and energetics (electron transport chain and oxidative phosphorylation). When we analyzed the functional role of the interactomes of Ppar-α/γ we noted that indeed both interactomes converged on lipid metabolism, mitochondrial bioenergetics and circadian processes (**Figure 3C**), all representing major cellular processes that are known to be dysregulated in IBD (23–30). These findings are consistent with the finding that *PPARA*, *PPARG* and *PPARGC1A* are located within clusters #1-2-3 and all of them are predicted to be progressively and simultaneously downregulated in IBD samples (**Figure 3D;** based on the IBD map, **Supplemental Figure 1**).

**Figure 3:**
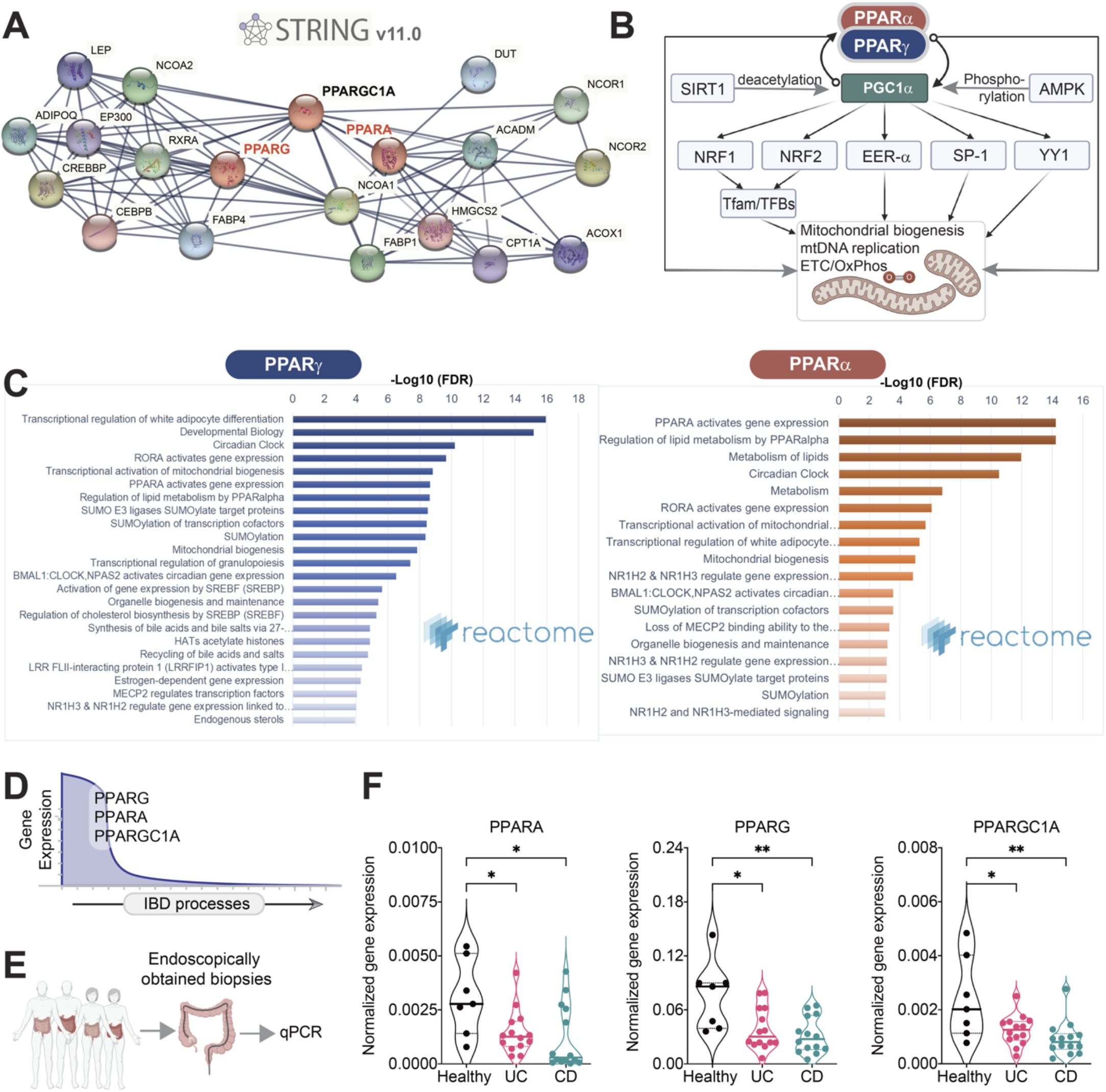
Rationalization of PPARα and PPARγ as targets in IBD. **(A)** A protein-protein interaction network (i.e., interactomes) for PPARα and PPARγ, generated using STRING v.11 (https://string-db.org). **(B)** Schematic summarizing the roles of PPARα, PPARγ and PGC1α on mitochondria biogenesis and function (based on). PGCl-α emerges as a critical hub for forward feedback loops. **(C)** Reactome pathway analyses (www.reactome.org) on PPAR-α and PPAR-γ interactomes in A show convergence on metabolism, mitochondria bioenergetics and the circadian clock. **(D)** Graphical visualization of the predicted changes in the expression of PPARA (PPAR-α), PPARG (PPAR-γ) and PPARGC1A (PGC1-α) genes during the progression of IBD processes (indicated with an arrow). **(E)** Schematic showing validation workflow; the expression of PPARA, PPARG and PPARGC1A transcript levels were assessed in the ileum/colon biopsies of IBD patients (UC=14 and CD= 14)) or healthy controls (n=7). **(F)** Violin plots display the qPCR results in E. Results are displayed as mean ± SEM. Significance was tested using one-way ANOVA followed by Tukey’s test for multiple comparisons. Significance: *, p < 0.05; **, p < 0.01.

### *PPARA, PPARG* and *PPARGC1A* are downregulated in Ulcerative colitis and Crohn’s Disease

Previous work demonstrated that both Ppar-α and Ppar-γ are highly expressed in the colon (31) and that their expression (proteins and mRNA) is downregulated (by ~60%) in active UC (32), in both inflamed and noninflamed areas (33). Moreover, the expression of Ppar-γ was significantly associated with disease activity (32). Polymorphisms have also been detected in Ppar-γ; while some studies found those to be associated with an increased risk for CD (34), others found no evidence suggesting any form of association with an increased disease risk (35). We collected endoscopically obtained biopsies from the colons of healthy (n = 7) and IBD (n = 14 and 14 of UC and CD, respectively) patients and assessed the levels of transcripts for *PPARA*, *PPARG* and *PPARGC1A* by qPCR (**Figure 3E**). We confirmed that all three transcripts were significantly downregulated in UC and CD samples compared to healthy; both *PPARG* and *PPARGC1a* were more significantly downregulated in CD compared to UC (**Figure 3F**). These findings are in keeping with the network-based predictions that these genes should be downregulated invariably in all IBD samples, regardless of disease subtype (see individual disease maps; **Supplemental Figure 4-5**). While both *PPARA* and *PPARG* are in cluster #2 in the UC map, *PPARG* and P*PARA* are in separate clusters, clusters 2 and 6, respectively, in the CD map (**Supplemental Figure 4-5**). Reactome pathway analyses implied that in the case of UC, the two nuclear receptors may co-regulate similar cellular homeostatic processes associated with cluster #2, i.e., mitochondrial biogenesis and translation initiation, infectious disease and detoxification of ROS (see **Supplemental Figure 4**). By contrast, in the case of CD, they may independently regulate diverse cellular processes that maintain cellular homeostasis; while *PPARG* is associated with cellular metabolism (TCA cycle) and inhibition of NFkB signaling, *PPARA* is associated with transcriptional activity of nuclear receptors, cholesterol biosynthesis and Met/Ras signaling (see **Supplemental Figure 5**). Taken together, these findings demonstrate that *PPARA/G* and *PPARGC1A* are downregulated in IBD and that they may regulate key pathophysiologic processes that are vital for cellular homeostasis. Findings support our AI-guided hypothesis that restoration of the expression of these genes will increase the expression of genes in C#1-2-3 and suppress the expression of, and that such increase.

### Synthesis and validation of PAR5359, a potent and specific Ppar-α/γ dual agonist

We next sought to identify appropriate pharmacologic tools to test our hypothesis. Direct agonism of *PPARGC1A*/Pgc1a was deemed as not feasible because the only known agonist, ZLN005, non-specifically and potently also activates AMPK (36), a target that is known to independently improve barrier integrity in IBD (9). Because Pgc1a is intricately regulated by feedback loops by Ppar-α/γ (**Figure 3B**), we strategized targeting Pgc1a indirectly via Ppar-α/γ instead. As for Ppar-α/γ dual agonists, we noted that all commercially available compounds lack ‘balanced’ agonistic activities (**Supplemental Table 3**) (37, 38). Drugs that have fallen aside due to safety concerns also lack balanced agonism; most of them are more potent on Ppar-γ than on Ppar-α by a log-fold (**Supplemental Table 3**). All these Ppar-α/γ dual agonists have been withdrawn due to safety concerns (22), but the cause of the ‘unsafe’ profile remains poorly understood. Saroglitazar, the drug that is the only active ongoing Phase-III trial (NCT03061721) in this class, has ~3 log-fold more potency on Ppar-a than Ppar-γ (39). Because our AI-guided approach suggested the use of simultaneous and balanced agonism, we favored the use of the only balanced and yet, specific Ppar-a/γ agonist described to date, PAR5359 (40, 41) (see **Supplemental Table 4**). In the absence of commercial sources or well-defined methods on how to synthesize this molecule, we generated PAR5359 in 4 synthetic steps (see details in *Methods*) and confirmed its specificity and the comparable agonistic activities using pure single Ppar-a [GW7647 (42)] or Ppar-γ [Pioglitazone (43)] agonists as controls (**Supplemental Figure 6**). With these potent and specific compounds as tools, and their doses adjusted to achieve the same potency, we set out to validate the network-based predictions using pre-clinical models.

### PAR5359 ameliorates *C. rodentium*-induced colitis, enhances bacterial clearance

We next sought to assess the efficacy of individual and dual agonists of our compounds in murine pre-clinical models. Ppar-α/γ’s role (or the role of their agonists) in protecting the gut barrier has been evaluated primarily in DSS-induced colitis (**Supplemental Table 1, 2**). This model is more related to the UC patient pathology. However, *BoNE* prioritized other models over DSS, many of which accurately recapitulate the Ppar-α and Pparγ-downregulation that is observed in the barrier-defect transcript signature in human IBD (**Figure 2E**). Among those, we chose *C. rodentium*-induced infectious colitis, a robust model to study mucosal immune responses in the gut and understand derailed host-pathogen interaction and dysbiosis, which is closely related with IBD, and more specifically, CD pathophysiology (44–46). This model is also known to emulate the bioenergetic dysbalance and mitochondrial dysfunction(47), both key cellular processes represented in C#1-2-3 within the IBD map (**Supplementary Figure 1**). Furthermore, this model requires the balanced action of macrophages (a cell line predicted to be the preferred cell type target; **Figure 2F**) to promote bacterial clearance and healing (48).

Colitis was induced by oral gavage of *C. rodentium* and mice were treated daily with the drugs *via* the intraperitoneal route (see **Figure 4A** for workflow; **Supplemental Figure 7A**). The dose for each drug was chosen based on their *EC_50_* on their respective targets so as to achieve equipotent agonistic activities (**Supplemental Figure 6**). Fecal pellets of individual mice were collected to determine the number of live bacteria present in the stool. As anticipated, the bacterial burden in all mice increased from day 5, reaching a peak on day 7, forming a plateau until day 11 before returning to pre-infection baseline by day 18 (**Figure 4B**). Compared against all other conditions, PAR5359-treated mice cleared the gut bacterial load significantly and rapidly (**Figure 4B-D**). *Citrobacter* infection was associated with significant epithelial damage and profuse infiltration of inflammatory cells and edema by day 7 (**Supplemental Figure 7B)** most of which resolved by day 18 (DMSO control; **Figure 4E**). Colons collected on day 7 showed that treatment with PAR5359 significantly reduced these findings when compared to vehicle (DMSO), PPARα and PPARγ agonists alone (**Supplemental Figure 7B)**. Unexpectedly, when we analyzed the colons on day 18, we noted persistent immune infiltrates in tissues in two treatment arms, pioglitazone and GW7647 (arrowheads; **Figure 4E**), but not in the vehicle control group, or those treated with PAR5359. These findings indicate that individual Ppar-α or Ppar-γ agonists may either retard bacterial clearance and/or induce an overzealous amount of inflammation, but the balanced dual agonist (PAR5359) may have effectively cleared infection and resolved inflammation. PAR5359 also reduced spleen inflammation as evidenced by a decreased spleen weight and length compared to vehicle control (**Supplemental Figure 7C-F**). The spleens of mice treated with DMSO, Ppar-α-alone agonist, GW7647 and Ppar-γ-alone agonist, Pioglitazone showed black-discoloration, presumably infarcts (arrows, **Supplemental Figure 7C, 7E**). Notably, the spleens of mice treated with Ppar-α-alone agonist, GW7647, showed a significant increase in spleen length (**Supplemental Figure 7D, 7F**).

**Figure 4:**
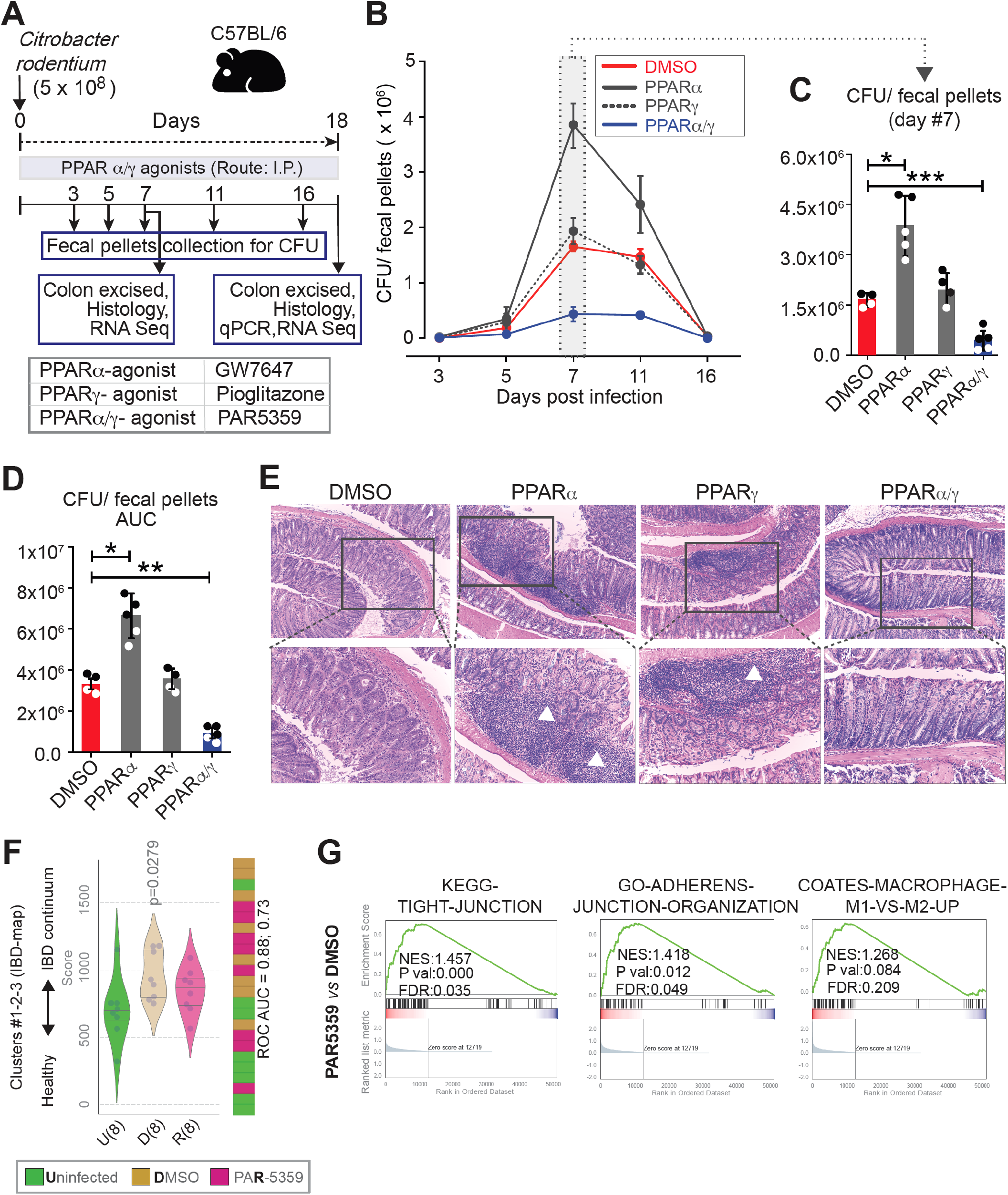
PPARα/γ dual agonists ameliorate *Citrobacter rodentium*-induced infectious colitis in mice. (**A**) Schematic summarizing the workflow for testing PPAR-targeted therapeutics in *C. rodentium*-induced colitis. Mice were gavaged with *C. rodentium* on day 0 and subsequently treated daily with PPAR agonists. Fecal pellets were collected to test viable bacterial burden, as determined by dilution plating and colony counting. Colons were excised on day 7 and 18 and analyzed using the indicated readouts. (**B-D**) Line graphs in B display time series of the burden of viable bacteria in feces. Scatter plots with bar graphs in C compare the peak burden of viable bacteria in feces on day 7. Scatter plots with bar graphs in D display the area under the curve (AUC) for the line graph in B. (**E**) Images display representative fields from H&E-stained colon tissues. Mag = 100x (top) and 200x (bottom). White arrowheads point to immune cell infiltrates. Statistics: All results are displayed as mean ± SEM. Significance was tested using two-way/one-way ANOVA followed by Tukey’s test for multiple comparisons. Significance: *, p < 0.05; **, p < 0.01, ***, p < 0.001. (**F**) Violin plots (left) display the deviation of expression of genes in Clusters #1-2-3 in the IBD network, as determined by RNA Seq on murine colons. Bar plot (right) displays the rank ordering of the samples. (**G**) Pre-ranked GSEA based on pairwise differential expression analyses (DMSO vs PAR5359 groups) are displayed as enrichment plots for epithelial tight (left) and adherens (middle) junction signatures and balanced macrophage processes (right). See also **Supplemental Figure 7** for the Day #7 results in the *C. rodentium*-induced colitis model, **Supplemental Figure 8** for extended GSEA analyses, and **Supplemental Figure 9** for the effect of PAR5359 on DSS-induced colitis in mice.

Taken together, these findings indicate that Ppar-α/γ dual agonist PAR5359 is superior in ameliorating *C. rodentium*-induced colitis than either Ppar-α or Ppar-γ agonist used alone. Treatment with the dual, but not the single agonists hastened bacterial clearance, resolved inflammation, and induced healing.

### PAR5359 resets the colonic gene expression changes induced by *C. rodentium* infection

Pharmacologic augmentation of *PPARA* and *PPARG* was hypothesized to be sufficient to upregulate genes in C#1-2-3, and restore the entire transcriptomic network to ‘healthy’ state *via* the invariant Boolean implication relationships between the genes/clusters. We asked if that was achieved. RNA sequencing (RNA-seq) studies were carried out on the *C. rodentium*-infected colons in each treatment group (**Figure 4A**). As expected, downregulation of genes in clusters #1-2-3 of the IBD-map was significant in infected untreated (DMSO control) *vs*. uninfected controls, indicative of network shift from health towards disease, and treatment with PAR5359 resisted such shift (**Figure 4F**).

Pre-ranked gene set enrichment analyses (GSEA) based on pair-wise differential expression analysis showed that when compared to DMSO control, dual Ppar-α/γ agonism with PAR5359, but not individual agonists Pioglitazone or GW7647 was able to significantly preserve epithelial junction signatures (both tight and adherens junctions) and balance macrophage processes (compare **Figure 4G** with **Supplemental Figure 8A**). These findings are in keeping with the predictions that epithelial cells and macrophages maybe the primary cell type of action for dual Ppar-α/γ agonists. Comparison of all treatment cohorts against each other revealed that although both PAR5359 and Pioglitazone were superior to GW7647 in maintaining some epithelial processes (differentiation, tight junctions) and macrophage processes (**Supplemental Figure 8B-E**), PAR5359 emerged as the only group that maintained homeostatic PPAR signaling in nature and extent as uninfected control (**Supplemental Figure 8F**).

Taken together, these findings suggest that dual agonists of Ppar-α/γ are sufficient to either resist network shift and/or reverse the disease network in the setting of colitis. They also offer clues suggestive of epithelial and macrophage processes, two key cellular components of innate immunity in the gut lining as major mechanisms. These transcriptome wide impacts suggest that Ppar-α/γ dual agonist PAR5359 is superior in restoring colon homeostasis in *C. rodentium*-induced colitis than either Ppar-α or Ppar-γ agonist used alone.

### PAR5359 ameliorates DSS-induced colitis

It is well known that no single mouse model recapitulates *all* the multifaceted complexities of IBD (49, 50). Because almost all studies evaluating Ppar-α/γ-modulators have been performed on the DSS-induced colitis model (**Supplemental Table 1-2**), we asked whether the Ppar-α/γ dual agonist PAR5359 can ameliorate colitis in this model. Mice receive intrarectal DMSO vehicle control or PAR5359 while receiving DSS in their drinking water (**Supplemental Figure 9A**). Disease severity parameters, i.e., weight loss, disease activity index, shortening of the colon and histology score were significantly ameliorated in the PAR5359-treated group (**Supplemental Figure 9A-E**). These findings show that the Ppar-α/γ-dual agonist, PAR5359, is also effective in DSS-induced colitis. It is noteworthy that the PAR5359 dual agonist offered protection in the DSS-model, because prior studies using the same model have demonstrated that Ppar-α agonists worsen (51, 52), and that the Ppar-γ agonists ameliorate colitis (53–55) (see **Supplemental Table 1-2**).

### PAR5359 promotes bacterial clearance with controlled production of ROS and inflammation in peritoneal macrophages

Next we sought to study the mechanism of action of PAR5359, and the target cell type responsible for the superiority of dual agonism over single agonism. Our AI-guided approach predicted crypt top epithelium and macrophages as site of action (**Figure 2F**). Based on prior studies with single agonists in cell-specific KO mice (**Supplemental Table 1**) and the phenotypes observed in our animal models (**Figure 4; Supplementary Figure 8-9**), single Ppar-γ agonism appears sufficient to protect the epithelium in chemical-induced colitis (dual agonism did not offer additional advantage). The advantage of dual agonism is apparent in the *Citrobacter*-colitis model, which most robustly recapitulates the paradoxical immune suppression in the setting of dysbiosis that is seen in IBD, and most prominently in CD (44, 56, 57). Because the intestinal macrophages, the known to be the initiators of immune response, are alternatively polarized in this model(58, 59), we hypothesized that balanced agonism may alter macrophage response to dysbiosis. To test this hypothesis, we incubated macrophages treated or not with the drugs and challenged them with CD-associated adherent invasive *E. coli (AIEC)-LF82*; this strain, originally isolated from a chronic ileal lesion from a CD patient (60). As for the source of macrophages, we isolated metabolically active primary murine peritoneal macrophages using Brewer thioglycolate medium using established protocols (61, 62). These macrophages are known to have high phagocytic activity (61) (**Figure 5A**). Thioglycolate-induced peritoneal macrophages (TG-PMs) were lysed, and viable intracellular bacteria were counted after plating on an agar plate. Pre-treatment with 1 μM PAR5359 and an equipotent amount of GW7647 (Ppar-α agonist) promoted bacterial clearance and reduced the bacterial burden when compared to vehicle control (**Figure 5B**). By contrast, pre-treatment with Pioglitazone (Ppar-γ agonist) inhibited bacterial clearance; notably, bacterial burden was significantly higher at both 3 h and 6 h after infection (**Figure 5B**). Reduced clearance of microbes in the latter was associated also with reduced cellular levels of reactive oxygen species (ROS) (**Figure 5C**); oxidative burst and induction of ROS is key component for effective bacterial killing (63, 64). PAR5359 did not interfere with the production of microbe-induced ROS, and the Ppar-α agonist (GW7647) was permissive to ROS induction (in fact, even induces it over bacteria-alone control) during initial time points after infection (**Figure 5C**).

**Figure 5:**
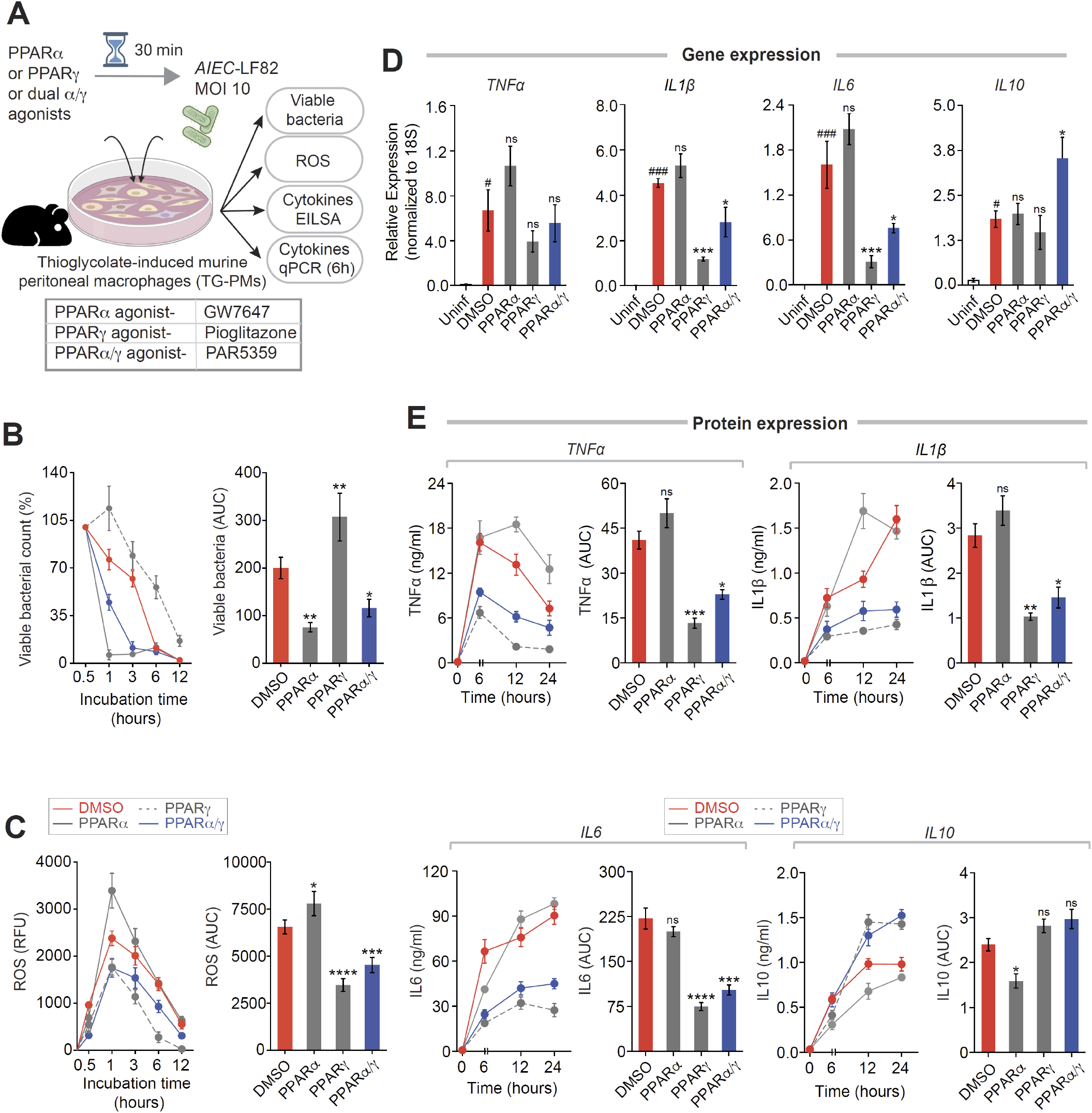
PPARα and PPARα/γ-dual agonists enhance, whereas PPARγ agonist delay bacterial (*AIEC*-LF82) clearance. (**A**) Schematic displays the experimental design and workflow. Thioglycolate-induced murine peritoneal macrophages (TG-PM) pretreated with PPAR agonists (see box, below; 20 nM GW7647, 10 μM Pioglitazone and 1 μM PAR5359) were infected with *AIEC-LF82 (MOI* 10) and subsequently analyzed for the bacterial count (Gentamicin protection assay), generation of cellular ROS, secretion of inflammatory cytokines (in supernatant media by ELISA) and the induction of cytokines (gene transcript analysis by qPCR). (**B**) Line graphs (left) display percent viable bacterial counts at indicated times after infection. Bar graphs (right) display the AUC. (**C**) Line graphs (left) and bar graphs (right) display the extent of ROS generation over time. (**D**) Bar graphs display the relative expression of transcripts of multiple cytokines (IL1β, IL6, TNFα and IL10). (**E**) Line graphs (left) and bar graphs (right) showing the levels of secreted cytokines in the media. Statistics: All results are from at least three independent experiments and results displayed as means ± SEM. Significance was tested using two-way/one-way ANOVA followed by Tukey’s test for multiple comparisons. Significance: ‘#’ significance over uninfected TG-PMs and ‘*’ shows significance over AIEC-LF82 infected cells. ns, non-significant, *, p < 0.05; **, p < 0.01, ***, p < 0.001, ****, p < 0.0001. See **Supplemental Figure 10** for similar bacterial clearance assays performed using *Salmonella enterica*.

These patterns of microbial clearance and cellular ROS were associated also with the expression of cytokines, as determined by qRT-PCR analyses (**Figure 5D**). As expected, infection of TG-PM with *AIEC*-LF82 induced *Il1β, Il6, Tnfα* and *Il10*. PAR5359 significantly and selectively suppressed the expression of the pro-inflammatory cytokines *Il1β, Il6* and *Tnfα* (but not the anti-inflammatory cytokine, *Il10*) (**Figure 5D**). By contrast, the Ppar-γ specific agonist pioglitazone significantly suppressed all the cytokines, while there was no effect of the Ppar-α specific agonist GW7647 (**Figure 5D**). ELISA studied on the supernatant media further confirmed these findings (**Figure 5E**), demonstrating that the effects in gene expression were also translated to the levels of secreted cytokine protein released by the macrophages in the supernatant.

It is noteworthy that for the most part, the qPCR (**Figure 5D**) and ELISA (**Figure 5E**) studies matched, except *Il10;* although pioglitazone appeared to suppress *Il10* mRNA, it did not suppress the levels of the IL10 protein, suggesting that Ppar-γ agonist is sufficient for an overall anti-inflammatory phenotype. Similarly, although GW7647 appeared to not affect *Il10* mRNA, it suppressed the levels of the IL10 protein, suggesting that Ppar-α agonist is sufficient for an overall pro-inflammatory phenotype. Similar findings were also observed in the case of another enteric pathogen, *S. enterica*, i.e., unlike the dual agonist, neither Ppar-α nor Ppar-γ agonist could enhance bacterial clearance with a modest induction of pro-inflammatory cytokines (significantly lower than control) and, yet, had no impact on anti-inflammatory IL10 production (**Supplemental Figure 10**).

Taken together, these results show that- (i) Ppar-γ agonism induces ‘tolerance’ by suppressing inflammation, inhibiting ROS production and delaying bacterial clearance; (ii) Ppar-α agonism enhances the induction of inflammation and ROS, and promotes bacterial clearance; and (iii) Ppar-α/γ dual agonism strikes a somewhat balanced response. The latter suppresses proinflammatory cytokines without suppressing anti-inflammatory cytokine *Il10*, and is permissive to inflammation and ROS induction that is optimal and sufficient to promote bacterial clearance.

### Ppar-α, but not Ppar-γ is required for the induction of inflammatory cytokines and ROS

To further dissect which nuclear receptors are responsible for the balanced actions of the dual agonist, we next used a set of highly specific and potent Ppar-α/γ inhibitors (**Supplemental Table 4**). We pre-treated TG-PMs with Ppar-α and Ppar-γ inhibitors, either alone, or in combination, followed by stimulation with bacterial cell wall component LPS (**Figure 6A**). As expected, LPS induced the cellular levels of ROS (**Figure 6B**) and inflammatory cytokines (**Figure 6C-D**) in TG-PMs significantly higher than in untreated control cells. Inhibition of Ppar-α suppressed the induction of cellular ROS and inflammatory cytokines, both at the level of gene and protein levels (**Figure 6B-D**). By contrast, inhibition of Ppar-γ did not interfere with either response (**Figure 6B-D**). Simultaneous inhibition of both Ppar-α and Ppar-γ mimicked the cellular phenotypes in the presence of Ppar-α inhibitors (**Figure 6B-D**), indicating that inhibition of Ppar-α is sufficient to recapitulate the phenotype of dual inhibition. Taken together, these findings indicate that Ppar-α is required for the proinflammatory response of macrophages.

**Figure 6:**
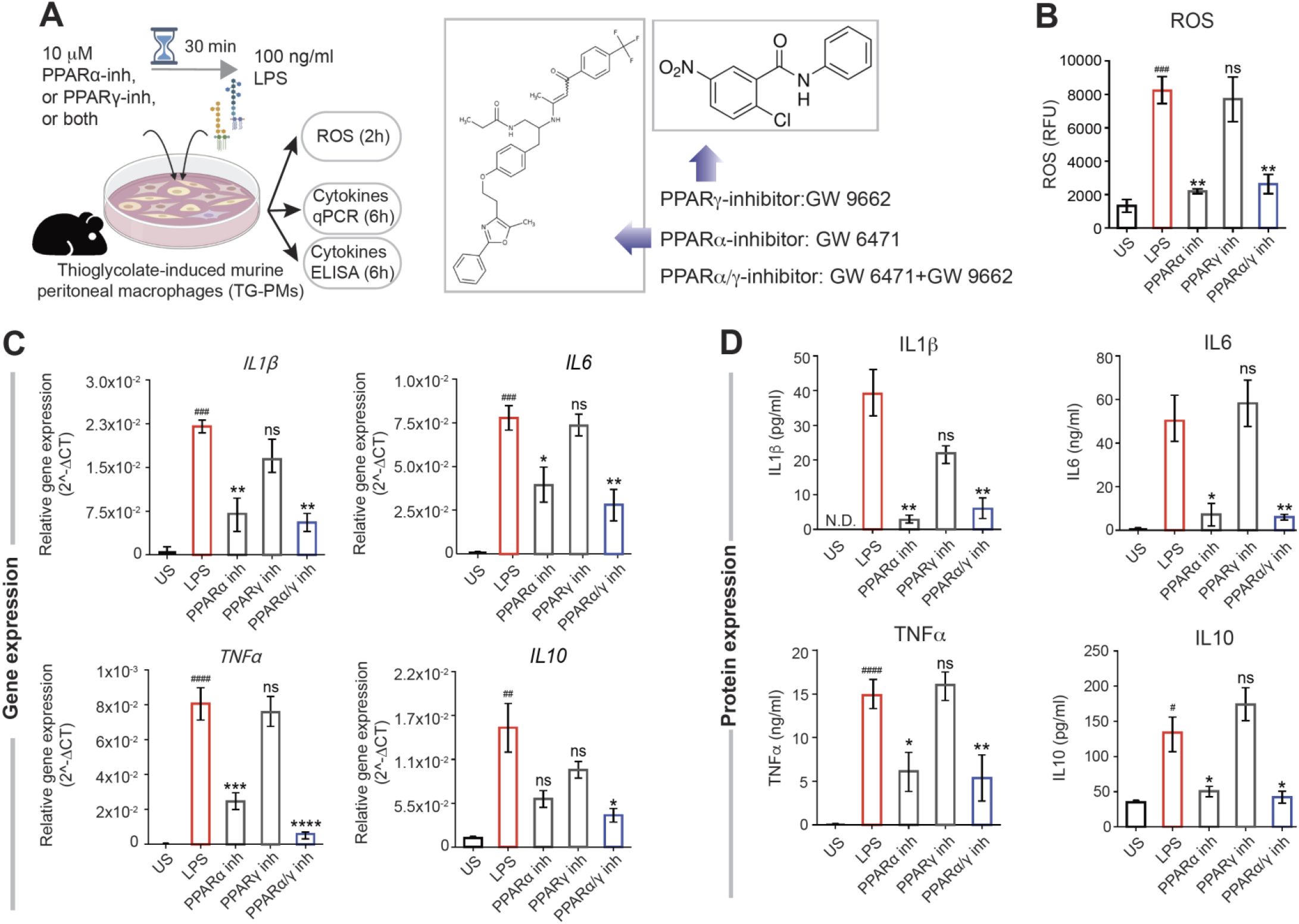
PPARa but not PPARγ is required for induction of cellular ROS and proinflammatory cytokines. **(A)** Schematic of experimental design. TG-PMs were pre-incubated with 10 μM PPARa or PPARγ inhibitors, either alone or in combination for 30 min prior to stimulation with 100 ng/ml LPS. Cells were analyzed at 2 and 6 h to estimate cellular ROS and cytokine induction, respectively. **(B-D)** Bar graphs display the levels of cellular ROS (B), relative levels of mRNA (C) and protein (D) expression of cytokines (IL1β, IL6, TNFα and IL10). Statistics: Results are from three independent experiments and displayed as mean ± SEM. One-way ANOVA followed by Tukey’s test for multiple comparisons was performed to test significance. Significance: ns: non-significant, *, p < 0.05; **, p < 0.01, ***, p < 0.001 and ****, p < 0.0001.

### Ppar-α/γ dual agonist PAR5359 promotes bacterial clearance in patient-derived PBMCs

In search of a pre-clinical human model for testing drug efficacy, we next assessed microbial handling by PBMCs derived from patients with IBD and compared them with that in age-matched healthy volunteers. We enrolled both male and female patients and both CD and UC (**Supplemental Table 5**). Consecutive patients presenting for routine care to the UC San Diego IBD clinic were enrolled into the study; the only exclusion criteria were failure to obtain informed consent for the study or active infections and/or disease flare. Peripheral blood collected in the clinic was freshly processed as outlined in **Figure 7A** to isolate PBMCs. Pre-treatment for 30 min with vehicle or PAR5359 was followed by infection for 1h. Subsequently, the cells were treated with gentamicin for 60 min to kill extracellular bacteria to assess intracellular bacterial burden at 1 and 6 h after the gentamicin wash.

**Figure 7:**
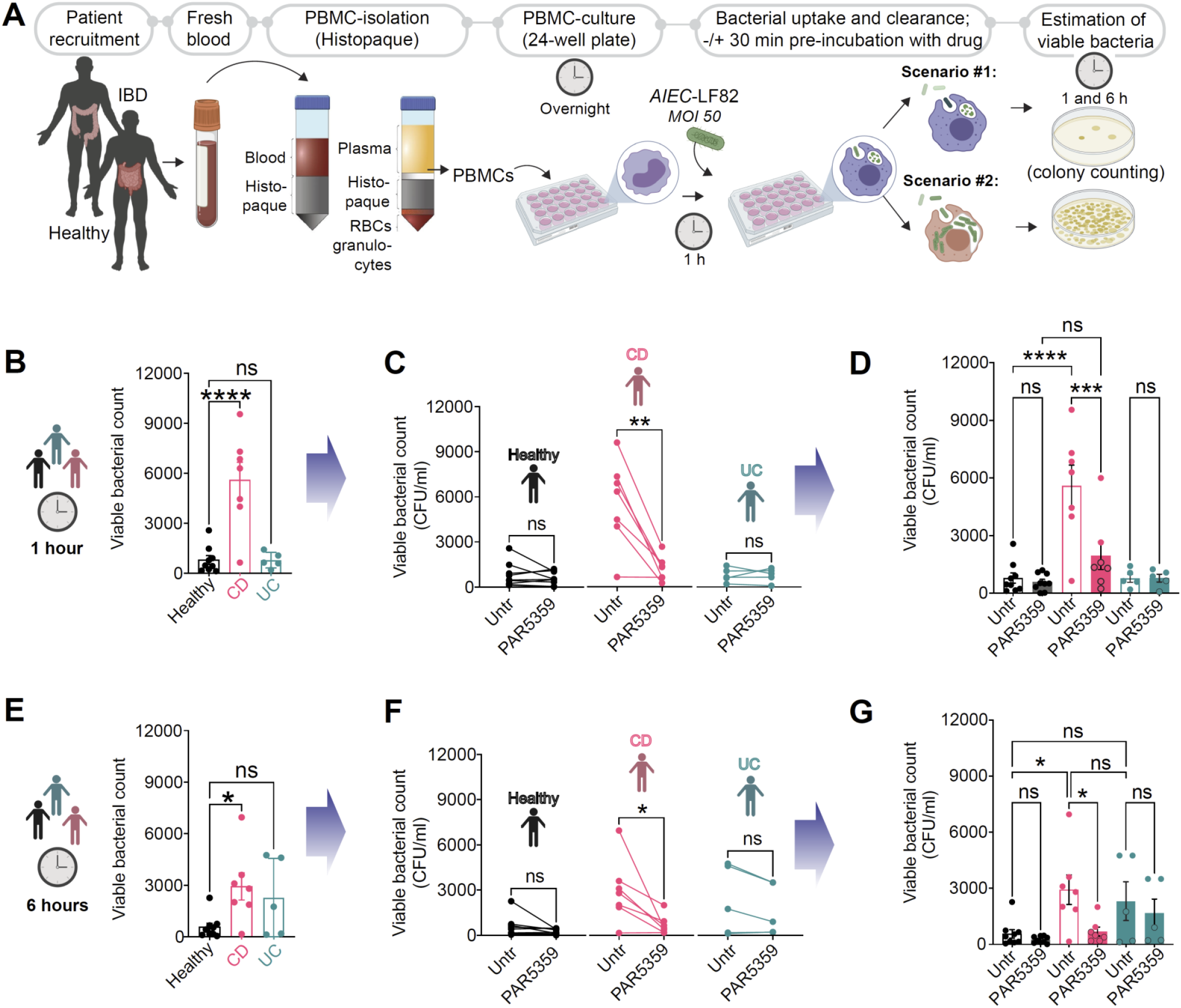
PPARα/γ dual agonist, PAR5359, promotes the clearance of *AIEC*-LF82 from CD patient-derived PBMCs. (**A**) Schematic displays the overall experimental design using human subjects (see **Supplemental Table 5** for patient demographics). Peripheral blood collected from healthy, CD and UC patients was used as a source of PBMCs. PBMCs were pre-treated for 30 min with 1 μM PPARα/γ agonists prior to infection with *AIEC-LF82 (MOI* 50) for 1h. PBMCs were subsequently treated with gentamicin to kill extracellular microbes for 60 min (~t0 h) prior to lysis and plating to determine the intracellular abundance of viable bacteria at t1h and t6h, as determined by dilution plating and colony counts (see *Methods* for details). Bar graphs with scatter plot display the abundance of viable intracellular bacteria at 1h (**B**) and 6h (**E**) after infection. Paired line plots display the rate of clearance of bacteria in individual subjects at 1h (**C**) and 6 h (**F**) after infection. Data in B-C of 1h infection is combined in (**D**) and data from E and F 6h infection is combined in (**G**) with statistics: Results are displayed as mean ± SEM (CD patient n=7, UC patients=6 and healthy n=9). Paired t-test or One-way ANOVA followed by Tukey’s test for multiple comparisons was performed to test significance. Significance: ns: non-significant, *, p < 0.05; **, p < 0.01, ***, p < 0.001 and ****, p < 0.0001.

Two observations were made: *First*, CD but not UC patient-derived PBMCs when infected with *AIEC-*LF82 showed an increased number of internalized viable bacteria when compared to healthy PBMCs (**Figure 7B, 7E**), indicative of either defective clearance and/or increased permissiveness to bacterial replication within the cells is limited to the CD. *Second*, pre-treatment with PAR5359 could improve clearance significantly (**Figure 7C-D, 7F-G**). These results indicate that bacterial clearance is delayed in PBMCs of patients with CD and that Ppar-α/γ dual agonism with PAR5359 can reverse that defect. The possibility that such reversal could be due to any direct bacteriostatic/-cidal effect of PAR5359 agonist was ruled out (see bacterial viability assay in **Supplemental Figure 11**). Our findings demonstrate that bacterial clearance is delayed primarily in CD and not UC are in keeping with the fact that delayed bacterial clearance from inflamed tissues (up to ~4-fold) is uniquely observed in CD (65). These findings are also in keeping with our own observation that the downregulation of *PPARG/PPARGC1A* was more prominent in patients with CD (**Figure 3E-F**). In fact, delayed clearance is one of the major reasons for persistent inflammation and disease progression among patients with CD (65, 66).

## DISCUSSION

Barrier-protection/restoration is the treatment endpoint for all clinical trials in IBD therapeutics; however, despite much success in the development of anti-inflammatory therapies (7, 67), barrier-protective therapeutics in IBD have been slow to emerge (68). Here we report the discovery of an effective barrier-protective therapeutic strategy in IBD identified using an AI-guided navigation framework (summarized in **Figure 8**). *First*, a network-based drug discovery approach (9) was used to identify, rationalize and validate dual and balanced agonism of Ppar-α/γ (but not one at a time) is necessary for therapeutic success. *Second*, we provided evidence in the form of proof-of-concept studies (in two different pre-clinical murine models) demonstrating that the simultaneous and balanced agonistic activation of the pair of PPARs as an effective barrier protective strategy in IBD. *Third*, we demonstrate that macrophages are one of the primary target cell type of this therapeutic strategy; dual agonist (but not single) was permissive to the induction of macrophage responses expected for optimal immunity without overzealous inflammation. There are *<u>three</u>* notable takeaways from this study, which are unexpected observations and/or insights that fill key knowledge gaps in the fields of – (a) network medicine, (b) IBD therapeutics and (c) macrophage biology.

**Figure 8:**
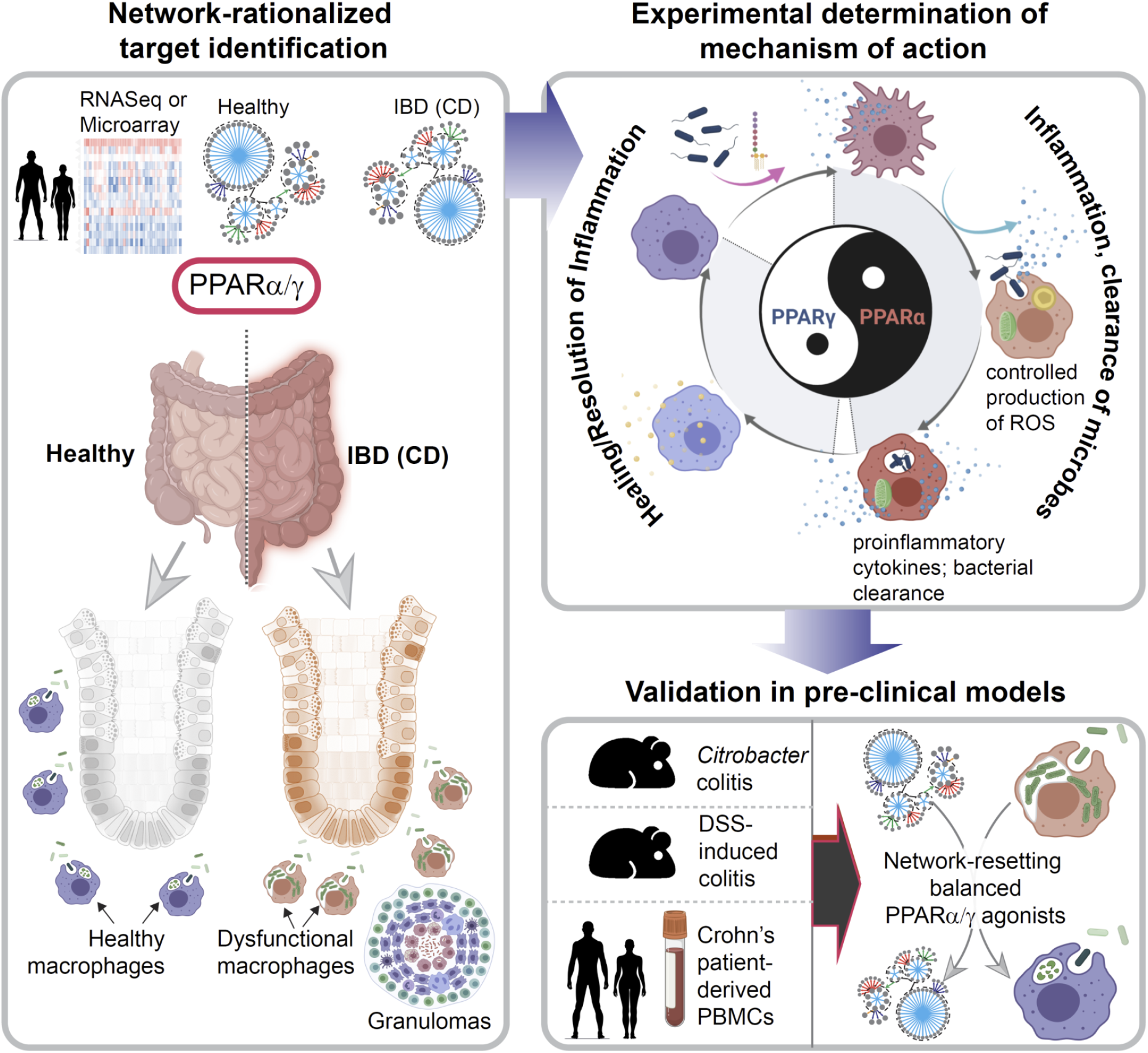
Summary of findings and working model. Schematic summarizes key approaches and findings of this study. First, network-rationalized target identification (*Left*) was performed using web-based platform that queries > 1000 IBD datasets [(9); see *Methods*] that served as ‘input’ to create a map of gene clusters that are progressively altered in the gut in the setting of IBD. Predictions are used to guide the choice of therapeutics (dual agonists of PPARa and PPARg that have a balanced agonistic potential for both PPARs), the choice of animal models of IBD, predict cell types of action (macrophage processes), and finally, the subtype of IBD that could benefit most based on the cell type of action (i.e., CD). Second, experimentally determined mechanism of action studies (*right, top*) showed that balanced actions of both PPARa and PPPARg enable the induction of bacterial clearance, resolution of inflammation and healing; PPARa is responsible for ROS and cytokine induction, whereas PPPARg is responsible for anti-inflammatory response and healing. The dual agonistic action was superior to each agonist used alone. Third, targets validation studies (*right, bottom*) in murine and human models confirm the use of PPARα/γ dual agonists for enhancing bacterial clearance and protection against colitis. When tested side-by-side in the infectious colitis model, the dual agonistic action was superior to each agonist used alone.

*First*, with regard to network medicine, the AI-guided approach we used here differs from the current practice in three fundamental ways: 1) Unlike most studies that prioritize targets based on Differential Expression Analysis (DEA, or integrated DEA) or Bayesian approaches, target identification and prediction, this work was guided by a Boolean implication network of continuum states in *human* disease (9); 2) Instead of conventional approaches of trial-and-error, intuitive guess and/or knowledge-based prioritization of study models (animal or cell-type of action), target validation in network-rationalized animal and cell-type models that most accurately recapitulate the role of the target(s) during disease progression; 3) Inclusion of human pre-clinical model (patient-derived PBMCs) for target validation, inspiring the concept of *Phase ‘0’ trials* that have the potential to personalize the choice of therapies. The combined synergy of these approaches validates a first-in-class macrophage modulator in addressing the broken gut barrier in IBD.

The impact of using such an approach is 4-fold: (i) Because the network approach used here relies on the fundamental invariant Boolean implication relationships between genes, and their patterns of changes in expression between healthy and IBD samples, such ‘rule of invariant’ implies that any given relationship and/or change in expression pattern annotated within the network *must* be fulfilled in every IBD patient. By that token, targets/drugs prioritized based on this network is expected to retain efficacy beyond inbred laboratory mice, into the heterogeneous patient cohorts in the clinic. (ii) This AI-guided approach not just helped compute pre-test probabilities of success (*“Therapeutic Index”*), but also helped pick models that are most insightful and appropriate to demonstrate therapeutic efficacy (e.g., *Citrobacter rodentium* infection-induced colitis) and to pinpoint the cell type and mechanism of action (microbial clearance by macrophages). This is noteworthy because the conventional approach in studying PPARs has been limited to the use of DSS-induced colitis (see **Supplemental Table 1-2**), which has often given conflicting results (see **Supplemental Table 2**). Ppar-γ agonists works best for the UC patients, perhaps because it is a potent inhibitor of proinflammatory cytokines and, as shown before, protects the intestinal epithelium(69). Our findings in the *Citrobacter* model imply that such single Ppar-γ agonism may worsen the macrophage dysfunction that is observed in the setting of CD, which is characterized by ineffective microbial clearance, insufficient proinflammatory response in the setting of luminal dysbiosis (28). In fact, without the use of the *Citrobacter rodentium* infectious colitis model, the deleterious effects of Ppar-γ agonists would have been overlooked. (iii) Having a computational framework improves precision in target identification; it is because of the emergence of the two PPARs (alongside their positive feedback regulator, Pgc1a) within our network, we rationalized their dual agonism as a preferred strategy (over single) and our experiments validated that prediction both *in vivo* and *in vitro*. This is noteworthy because conventional approaches have demonstrated a protective role of Ppar-γ agonists and a conflicting (both protective and exacerbating) role of Ppar-α in IBD (52, 70–72); the advantage of dual agonism has neither been rationalized nor tested. (iv) The ‘target report card’, like the one shown here, is a project navigation tool that is geared to streamline decision-making (i.e., which genes, which animal models, which cell type/cellular process, what is the likelihood of success, etc.), which in turn should reduce attrition rates, waste and delays; the latter are well-recognized flaws in the current process of drug discovery.

*Second*, regarding IBD therapeutics, our studies demonstrate that single or unbalanced combinations of Ppar agonists are inferior to dual/balanced agonists. Conventional and reductionist approaches have inspired numerous studies with single Ppar agonists over the past decade (**Supplemental Tables 1-2**). However, given the devastating side effects of most single or unbalanced Ppar-α/γ agonists (**Supplemental Table 3**), translating to the clinic beyond a Phase II trial (17, 73, 74) has not been realized. Because the therapeutic index for the dual Ppar-α/γ agonists matches that of other FDA-approved targets/drugs, it is predicted that barring unexpected side effects, dual Ppar agonists are likely to be effective as barrier-protective agents. As for side effects, we noted is that balanced Ppar-α/γ agonists are rare; while all dual Ppar-α/γ agonists that have been discontinued due to side effects happen to be either single (only Ppar-γ) or ‘unbalanced’ (Ppar-γ >> Ppar-α agonistic activity), the newer generation formulations that are currently in the clinical trial have a reversed agonistic potency (Ppar-α >> Ppar-γ agonistic activity) (see **Supplemental Table 3**). Because macrophage responses require finetuning (discussed below), our studies show how unopposed agonism of either Ppar-γ or Ppar-α is harmful and can impair/dysregulate the way macrophages respond when microbes breach past the gut barrier. It is possible that many of the side effects of the discontinued thiazolidinediones are due to their inability to achieve that ‘optimal’ spectrum of macrophage function.

*Third*, when it comes to macrophage biology, this work sheds some unexpected and previously unforeseen insights into the role of the PPARs in the regulation of macrophage processes. Extensively studied for over ~3 decades, PPARs are known to regulate macrophage activation in health and disease (75). Targeting PPARs as a host-directed treatment approach to infectious/inflammatory diseases appears to be a sound strategy because they regulate macrophage lipid metabolism, cholesterol efflux, inflammatory responses (ROS and cytokine production), apoptosis, and production of antimicrobial byproducts (76). We found that unopposed Ppar-γ activation suppresses bacterial clearance and blunts the induction of proinflammatory (but not anti-inflammatory, IL10) cytokines and ROS in response to infection both *in vivo* and *in vitro*. In other words, and consistent with prior reports, Ppar-γ activation suppressed inflammation at the cost of impairing immunity. Our findings are in keeping with the findings of a systematic review and meta-analysis of 13 long-term randomized controlled trials that involved 17,627 participants (8,163 receiving Ppar-γ agonists and 9,464 receiving control drugs)(77). Long-term (~1–5.5 y) use of Ppar-γ agonists increases the risk of pneumonia or lower respiratory tract infection significantly, some of which result in hospitalization, disability, or death(77). In the case of Ppar-α, unopposed activation-induced ROS and proinflammatory cytokines and accelerated bacterial clearance. Inhibitor studies further confirmed that Ppar-α was required for these responses (**Figure 6**). These findings are in keeping with others’ showing that Ppar-α, but not Ppar-γ is required for NADPH-induced ROS formation both in human and murine macrophages (78). Ppar-α agonists induce the expression of NADPH oxidase subunits p47(phox), p67phox, and gp91phox, which are all essential functional components of NADPH complex (78). Dual and balanced Ppar-α/γ agonism enhanced bacterial clearance with only a moderate induction of proinflammatory cytokines or ROS. Such a response ensures that the macrophage functions within a ‘goldilocks’ zone, mounting inflammation that is just sufficient for microbial clearance and immunity. In our analysis, the only other PPAR-related gene within the IBD network, i.e., Pgc1a, and its role within the Ppar-α/γ axis suggests that the intricate network of forward feedback loops orchestrated by Pgc1a may be critical for achieving the critical balance between immunity and inflammation, which is a key outcome of the dual Ppar-α/γ agonists.

Because previous studies using cell-specific gene depletion have indicated that the barrier-protective role of Ppar-γ may be mediated *via* cells other than the macrophages (54), namely, the T cells (79) and the epithelial cells (80), it is possible that the dual Ppar-α/γ agonists also act on those cells, promoting bacterial clearance and balancing cellular bioenergetics, ROS and cytokine production, in manners similar to that we observe in macrophages.

Taken together, our study uses an unconventional approach to rationalize and validate the use of Ppar-α/γ dual agonists as first-in-class barrier protective macrophage modulators in the management of IBD. The approach is powerful because it leverages the precision of mathematics (*Boolean* algebra of *logic*) and the fundamental invariant patterns in gene expression (Boolean Implications). The AI-navigated drug discovery approach defined here could serve as a blueprint for future studies not just in IBD, but in any other such complex chronic diseases.

## AUTHOR CONTRIBUTIONS

P. G., D.S., S.D. and G.D.K conceptualized, supervised, administered the project and acquired funding to support it. G.D.K., I.M.S., M.S.A, E.V., F.U., J.R.S were involved in data curation and formal analysis. D.S. developed computational modeling and software for analysis and performed all computational analysis. G.D.K., I.M.S., M.S.A, E.V., F.U., conducted animal studies for colitis. G.D.K., I.M.S., M.S.A, E.V., F.U., and S.D., performed cell and tissue analysis including qPCR, ROS, bacterial clearance, ELISA. D.T. assisted G.D.K. for qPCR and analysis. J.R.S., G.L. and J.Y. prepared PAR5359 compound. W.J.S. provided key resources for human subjects. D.S., G.D.K. and P.G. prepared figures for data visualization, wrote original draft, reviewed and edited the manuscript. All co-authors approved the final version of the manuscript.

## ACKNOWLEDGMENTS

We thank Dharanidhar Dang (UCSD) for comments and critiques during the preparation of the manuscript. This work was supported by National Institutes for Health (NIH) grants R01-AI141630 (to PG), DK107585 (to SD). PG, SD, and DS were also supported by the Leona M. and Harry B. Helmsley Charitable Trust and the NIH (UG3TR003355, UG3TR002968 and R01-AI55696). GDK was supported through The American Association of Immunologists Intersect Fellowship Program for Computational Scientists and Immunologists. J.S. acknowledges support from the Interfaces Training Grant at UCSD (NIH T32EB009380). Authors thank to Lee Swanson, Courtney Tindle, Stella-Rita Ibeawuchi, Julian Tam and Madhubanti Mullick for their comments, feedback and technical support. This manuscript includes data generated at the UC San Diego Institute of Genomic Medicine (IGC) using an Illumina NovaSeq 6000 that was purchased with funding from a National Institutes of Health SIG grant (#S10 OD026929). Additionally, a P30 grant (NIH/NIDDK, P30DK120515) subsidized the RNA Seq and histology work showcased here.

