## Supplementary material for "Artificial Intelligence-rationalized balanced PPARα/γ dual agonism resets the dysregulated macrophage processes in inflammatory bowel disease": Methods, Supplemetal Figures S1-S11, Supplemental Table 1-5



**Conflict of interest statement:** S.D, D.S and P.G have a patent on the methodology. Barring this, the authors have declared that no conflict of interest exists.


**Runnning title:** AI-guided macrophage modulation in IBD


***Correspondence to:**

**William J. Sandborn, M.D.;** Professor, Department of Medicine, University of California San Diego; 9500 Gilman Drive, MC 0956, La Jolla, CA 92093-0831.

**Soumita Das, Ph.D.;** Associate Professor, Department of Pathology, University of California, San Diego; 9500 Gilman Drive, George E. Palade Bldg, Rm 256, 239; La Jolla, CA 92093.

**Debashis Sahoo, Ph.D;** Assistant Professor, Department of Pediatrics, University of California San Diego; 9500 Gilman Drive, MC 0730, Leichtag Building 132; La Jolla, CA 92093-0831.

**Pradipta Ghosh, M.D.;** Professor, Departments of Medicine, and Cell and Molecular Medicine, University of California San Diego; 9500 Gilman Drive (MC 0651), George E. Palade Bldg, Rm 232, 239; La Jolla, CA 92093. Phone: 858-822-7633: Fax: 858-822-7636:

#### CATALOG OF SUPPLEMENTARY MATERIALS


1. *Methods (Page 3-14)*
2. *Table of Resources (Page 15-16)*
3. *Supplemental Figures and Legends (S1-S10, Page 17-27)*
4. *Supplemental Tables (1-5, Page 28-32)*
5. *Supplementary Bibliography (Page 33-36)*

### METHODS

#### *Computational methods*

##### A Boolean network map of IBD

A Boolean implication network (BIN) was created earlier [(1); **Supplemental Figure 1A**], and this network is comprised of clusters of genes, interconnected by Boolean Implication Relationships (BIRs). The concepts, mathematical, statistical, datasets that went into building this map is detailed in (1), and briefly mentioned here.

##### Gene expression databases

Publicly available human colon tissue gene expression databases were downloaded from the National Center for Biotechnology Information (NCBI) Gene Expression Omnibus website (GEO) (2-4). If the dataset is not normalized, RMA (Robust Multichip Average)(5, 6) is used for microarrays and TPM (Transcripts Per Millions)(7, 8) is used for RNASeq data for normalization. We used log2(TPM+1) to compute the final log-reduced expression values for RNASeq data. Accession numbers for these crowdsourced datasets are provided in the figures and manuscript. All of the above datasets were processed using the Hegemon data analysis framework (9-11).

##### Boolean Analysis

*Boolean logic* is a simple mathematic relationship of two values, i.e., high/low, 1/0, or positive/negative. The Boolean analysis of gene expression data requires first the conversion of expression levels into two possible values. The *StepMiner* algorithm is reused to perform Boolean analysis of gene expression data (12). The *Boolean analysis* is a statistical approach that creates binary logical inferences that explain the relationships between phenomena. Boolean analysis is performed to determine the relationship between the expression levels of pairs of genes. The *StepMiner* algorithm is applied to gene expression levels to convert them into Boolean values (high and low). In this algorithm, first the expression values are sorted from low to high and a rising step function is fitted to the series to identify the threshold. Middle of the step is used as the *StepMiner* threshold. This threshold is used to convert gene expression values into Boolean values. A noise margin of 2-fold change is applied around the threshold to determine intermediate values, and these values are ignored during Boolean analysis. In a scatter plot, there are four possible quadrants based on Boolean values (**Supplemental Figure 2A**): (low, low), (low, high), (high, low), (high, high).

##### Invariant Boolean implication relationships

A Boolean implication relationship is observed if any one of the four possible quadrants or two diagonally opposite quadrants are sparsely populated. Based on this rule, there are six different kinds of Boolean implication relationships. Two of them are symmetric: equivalent (corresponding to the highly positively correlated genes), opposite (corresponding to the highly negatively correlated genes). Four of the Boolean relationships are asymmetric, and each corresponds to one sparse quadrant: (low => low), (high => low), (low => high), (high => high). BooleanNet statistics (Equations listed below) is used to assess the sparsity of a quadrant and the significance of the Boolean implication relationships (12, 13). Given a pair of genes A and B, four quadrants are identified by using the StepMiner thresholds on A and B by ignoring the Intermediate values defined by the noise margin of 2-fold change (+/- 0.5 around StepMiner threshold). The number of samples in each quadrant is defined as $a_{00}$, $a_{01}$, $a_{10}$, and $a_{11}$ (**Supplemental Figure 2A**). The total number of samples where gene expression values for A and B are low is computed using the following equations.

$$nA_{low} = (a_{00} + a_{01}), nB_{low} = (a_{00} + a_{10}),$$

Total number of samples considered is computed using following equation.

$$total = a_{00} + a_{01} + a_{10} + a_{11}$$

Expected number of samples in each quadrant is computed by assuming independence between A and B. For example, expected number of samples in the bottom left quadrant $e_{00} = \hat{n}$ is computed as probability of A low ($(a_{00} + a_{01})$/total) multiplied by probability of B low ($(a_{00} + a_{10})$/total) multiplied by total number of samples. Following equation is used to compute the expected number of samples.

$$n = a_{ij}, \hat{n} = (nA_{low}/total * nB_{low}/total) * total$$

To check whether a quadrant is sparse, a statistical test for ($e_{00} > a_{00}$) or ($\hat{n} > n$) is performed by computing $S_{00}$ and $p_{00}$ using following equations. A quadrant is considered sparse if $S_{00}$ is high ($\hat{n} > n$) and $p_{00}$ is small.

$$S_{ij} = \frac{\hat{n} - n}{\sqrt{\hat{n}}}$$

$$p_{00} = \frac{1}{2}\left(\frac{a_{00}}{(a_{00} + a_{01})} + \frac{a_{00}}{(a_{00} + a_{10})}\right)$$

A threshold of $S_{00} >$ sthr and $p_{00} <$ pthr to check sparse quadrant. A Boolean implication relationship is identified when a sparse quadrant is discovered using following equation.

***Boolean Implication*** = ($S_{ij} >$ sthr, $p_{ij} <$ pthr)

A relationship is called Boolean equivalent if top-left and bottom-right quadrants are sparse (**Supplemental Figure 2B**).

$$Equivalent = (S_{01} > sthr,\ P_{01} < pthr, S_{10} > sthr, P_{10} < pthr)$$

Boolean opposite relationships have sparse top-right ($a_{11}$) and bottom-left ($a_{00}$) quadrants.

$$Opposite = (S_{00} > sthr,\ P_{00} < pthr, S_{11} > sthr, P_{11} < pthr)$$

Boolean equivalent and opposite are symmetric relationship because the relationship from A to B is same as from B to A. Asymmetric relationship forms when there is only one quadrant sparse (A low => B low: top-left; A low => B high: bottom-left; A high=> B high: bottom-right; A high => B low: top-right). These relationships are asymmetric because the relationship from A to B is different from B to A. For example, A low => B low and B low => A low are two different relationships.

A low => B high is discovered if bottom-left ($a_{00}$) quadrant is sparse and this relationship satisfies following conditions.

$$A\ low => B\ high = (S_{00} > sthr,\ P_{00} < pthr)$$

Similarly, A low => B low is identified if top-left ($a_{01}$) quadrant is sparse.

$$A\ low => B\ low = (S_{01} > sthr,\ P_{01} < pthr)$$

A high => B high Boolean implication is established if bottom-right ($a_{10}$) quadrant is sparse as described below.

$$A\ high => B\ high = (S_{10} > sthr,\ P_{10} < pthr)$$

Boolean implication A high => B low is found if top-right ($a_{11}$) quadrant is sparse using following equation.

$$A\ high => B\ low = (S_{11} > sthr,\ P_{11} < pthr)$$

For each quadrant, a statistic $S_{ij}$ and an error rate $p_{ij}$ is computed. $S_{ij} > 2.5$ and $p_{ij} < 0.1$ are the thresholds used on the BooleanNet statistics to identify Boolean implication relationships (BIRs). False discovery rate is computed by randomly shuffling each gene and computing the ratio of the number of Boolean implication relationship discovered in the randomized dataset and original dataset. For IBD dataset the false discovery rate was less than 0.001.

Boolean Implication analysis looks for invariant relationship across all the different types of samples regardless of the conditions and treatment protocols. Therefore, it does not distinguish the sample types when discovering Boolean implication relationships (**Supplemental Figure 2C-D**). We assume that there are fundamental invariant Boolean implication formula that are satisfied by every sample regardless of their type (in this context it is limited to healthy and IBD colonic biopsies including both UC and CD). This means normal, UC and CD samples share the same fundamental relationships.

**Inflammatory bowel disease (IBD) datasets**

Both Peters-2017 GSE83687 and Arijs-2018 GSE73661 dataset were indepndently prepared for Boolean analysis by filtering genes that have reasonable dynamic range of expression values by analyzing the fraction of high and low values identified by the StepMiner algorithm (14). Any probeset or genes that contain less than 5% of high or low values or do not have a big dynamic range are dropped from the analysis (for Peters-2017 dataset 7659/23228 genes dropped - 33%). To check if pairwise Boolean implication relationships are consistent between two datasets, every gene in Peters-2017 dataset is mapped to the best probeset (identified by the biggest dynamic range) in the Arijs-2018 dataset, and genes/probesets that do not match are dropped from the analysis (4841/23228 genes dropped – 21%). Finally, 44% (10232/23228) of genes were not used in the Boolean Implication Network because their expression did not have a sufficient range. Since RNA-Seq expression values have slightly different characteristics than microarray expression values, the consistency of Boolean implication relationship was determined by using BooleanNet statistics in both datasets and a Pearson's correlation coefficient in the Arijs-2018 dataset. A Pearson's correlation coefficient > 0.5 was considered compatible with Equivalent, High => High, and Low => Low Boolean implication relationships. Similarly, a Pearson's correlation coefficient < -0.25 was considered compatible with Opposite, High => Low, and Low => High Boolean implication relationships. The Boolean model is tested in several human datasets, each comprised of a heterogeneous collection of samples to demonstrate reproducibility (GSE16879, GSE59071, GSE48958, GSE50594, GSE37283, E-MTAB-7604). We have collected publicly available gene expression datasets derived from mouse models of IBD (DSS bulk GSE42768, DSS epithelium E-MTAB-5249, TNBS GSE53835, Citrobacter GSE90577, adoptive T-cell transfer ACT GSE87317, ACT GSE27302, IL10 -/- GSE39859, TNFR1 -/- GSE107933, TNFR2 -/- GSE65408) to test whether human Boolean models performs well in mice. The gene name conversion from human to mouse is performed using human genome GRCh38.95 ensembl IDs and mapping data exported from ensemble BioMart web-interface.

**Generation of Target report card**

A target report card is generated for one target or multiple targets to predict the efficacy of a potential drug. The target report card contains five different sections as described below: (1) Therapeutic index, (2) IBD outcome, (3) Network-prioritized mouse model, (4) estimation of gender bias, (5) Predicted tissue cell type of action.

**Target report card – Therapeutic index**

Therapeutic index is a number (lower the better) assigned to one target or multiple targets that predicts the efficacy of a potential drug. Therapeutic index is computed by measuring the strength of Boolean implication relationship with PRKAB1 (**Supplemental Figure 2E**). Since PRKAB1 is an agonist, if $S_{11} > 0$ then gene A is an Antagonist because top-right quadrant will have fewer samples than expected and PRKAB1 high will be associated with gene A low. For antagonist, top-right and bottom-left quadrants are expected to be sparse. Therefore, Tindex for antagonist is computed as follows:

$$T_{index} = \frac{1}{4}\left(\frac{0.3}{(S_{00} + 1)} + \frac{0.3}{(S_{11} + 1)} + p_{00} + p_{11}\right)$$

Similarly, if $S_{11} > 0$ then gene A is an Agonist because top-right quadrant will have more samples than expected and PRKAB1 high will be associated with gene A high. For agonist, top-left and bottom-right quadrants are expected to be sparse. Therefore, Tindex for agonist is computed as follows:

$$T_{index} = \frac{1}{4}\left(\frac{0.3}{(S_{01} + 1)} + \frac{0.3}{(S_{10} + 1)} + p_{01} + p_{10}\right)$$

Therapeutic indices range from 0.36 to 0.027 where the most effective drug targets will be close to 0.027 and abandoned drug targets will be close to 0.36. Since all the currently known FDA approved drug targets have therapeutic indices less than 0.1, we set this number as a threshold to identify effective drug targets. Lower therapeutic indices means stronger Boolean Implication relationship with PRKAB1 which predicted phase III successes for many drugs in IBD (1). Only four out of 16 targets have therapeutic indices less than 0.1. For effectiveness, we also check EMT and Inflammation scores in addition to the therapeutic index. Effective targets are observed to have better scores for both EMT and Inflammation, and they are likely to be present in both EMT and Inflammation Boolean paths. See section "**Identification of Epithelial-Mesenchymal and Inflammation-Fibrosis continuum**".

**Target report card – IBD outcome**

Several datasets with annotations of IBD (normal vs IBD, GSE73661, GSE16879, GSE59071, GSE48958) as well as the aggressiveness of IBD such as active from inactive disease (GSE59071, GSE48958), responders from non-responders receiving two different biologics, Infliximab or Vedolizumab (GSE73661, GSE16879, GSE50594, E-MTAB-7604), and even distinguished those with the quiescent disease with or without remote neoplasia (GSE37283) were used to assess the strength of association of drug targets with IBD outcome. See section "**Generation of heat maps and drug targets score**" of how drug targets to score is computed, samples are ordered and association with disease outcome is measured. A list of barplots is used to visualize the sample ordering and the association with disease outcome.

**Target report card – Network-prioritized mouse model**

Drug targets score is computed for the mouse model IBD datasets (DSS bulk GSE42768, DSS epithelium E-MTAB-5249, TNBS GSE53835, Citrobacter GSE90577, adoptive T-cell transfer ACT GSE87317, ACT GSE27302, IL10 -/- GSE39859, TNFR1 -/- GSE107933, TNFR2 -/- GSE65408) to test how combined gene expression values of the drug targets are associated with disease annotation. See section "**Generation of heat maps and drug targets score**" of how drug targets score is computed, samples are ordered and association with disease outcome is measured. A list of barplots is used to visualize the sample ordering and the association with disease outcome.

**Target report card – estimation of gender bias**

A box plot of the gene expression values of the individual target gene is computed in the Peters-2017 GSE83687 dataset to test if there are significant gender-associated differences. The box plots of individual genes for both males and females are plotted side-by-side to visualize the differences.

**Target report card – Predicted tissue cell type of action**

It is important to know which cell types are relevant for the optimal action of drug targets. We assume that the drug action is dictated by cell type-specific expression patterns of the drug targets. We predict cell type-specific expression patterns using various techniques including correlation, standard deviation, and previously published MiDReG algorithm (13). We assembled several gene expression databases for this task. A large human colon tissue database (n=1911) was assembled by pooling several normal colon, adenoma, and colorectal cancer datasets from NCBI GEO (**Supplemental Figure 3A**). All the samples in this database were analyzed using bulk tissue in Affymetrix U133 Plus 2.0 microarray platform. A large human colorectal cancer cell line database (n = 264) was prepared to identify genes expressed in epithelium because these are likely homogeneous and devoid of stromal tissue such as fibroblasts and immune cells. Microarray datasets of FACS purified macrophages, FACS purified GI fibroblasts, and FACS purified lymphocytes were downloaded using GSE134312, GSE63626, and GSE24759, respectively (**Supplemental Figure 3A**). The algorithm that predicts whether a gene is expressed in top, bottom, lymphocytes, macrophages, and fibroblasts is described in a flow chart (**Supplemental Figure 3B**). MiDReG algorithm is performed on the human colon tissue database (n=1911) to predict top/bottom of the crypt marker (9, 13). Boolean implication "KRT20 low => X low" is used to predict the expression of gene X at the top of crypt (9). Boolean implication "KRT20 low => X high" is used to predict the expression of gene X at the bottom of the crypt. Since the human colon tissue database (n=1911) contains bulk tissue samples, expression of gene X is restricted to the epithelium by filtering the gene expression in the human colorectal cancer cell line database (n = 264). LGR5 correlation > 0.8 is performed in the FACS purified colon crypt dataset (GSE31255) to predict bottom of the crypt markers independently. Standard deviation > 0.5 in human B cells and T cells (GSE24759) is used to predict lymphocyte-specific expression. Genes expressed in human macrophages and fibroblast is predicted by computing the StepMiner threshold on the bulk datasets GSE134312 and GSE63626, respectively, which is compared to the StepMiner threshold obtained in the original human global tissue dataset (GSE119087, n = 25,955). While both PPARG and PPARA are predicted to be expressed in top of the crypt and macrophages, PPARG is predicted to be expressed in fibroblasts in addition (**Supplemental Figure 3C**).

**Construction of a Network of Boolean Implications**

A Boolean implication network (BIN) is created by identifying all significant pairwise Boolean implication relationships (BIRs) that are consistent in both Peters-2017 GSE83687 and Arijs-2018 GSE73661 datasets independently (**Supplemental Figure 1A**) (15, 16). The Boolean implication network contains the six possible Boolean relationships between genes in the form of a directed graph with nodes as genes and edges as the Boolean relationship between the genes. The nodes in the BIN are genes and the edges correspond to BIRs. Equivalent and Opposite relationships are denoted by undirected edges and the other four types (low => low; high => low; low => high; high => high) of BIRs are denoted by having a directed edge between them. The network of equivalences seems to follow a scale-free trend; however, other as we generated PAR5359 through 2 intermediate steps, symmetric relations in the network do not follow scale-free properties. BIR is strong and robust when the sample sizes are usually more than 200 (from our experience of using Boolean Implication for more than 10 years). All our previous papers use thousands of diverse samples to establish Boolean implication relationships. Boolean Implication analysis is carried out for the first time in such low number of samples such as the selected IBD GSE83687 dataset (n = 134). We have demonstrated that we have a reasonable False Discovery Rate (< 0.001) when S > 2.5 and p < 0.1 are used. The IBD dataset was prepared for Boolean analysis by filtering genes that had a reasonable dynamic range of expression values. When the dynamic range of expression values was small, it was difficult to distinguish if the values were all low or all high or there were some high and some low values. Thus, it was determined to be best to ignore them during Boolean analysis. The filtering step was performed by analyzing the fraction of high and low values identified by the StepMiner algorithm (14). Any probe set or genes which contained less than 5% of high or low values were dropped from the analysis.

**Generation of Clustered Boolean Implication network**

Clustering was performed in the Boolean implication network to dramatically reduce the complexity of the network (**Supplemental Figure 1B**). A clustered Boolean implication network (CBIN) was created by clustering nodes in the original BIN by following the equivalent BIRs. One approach is to build connected components in a undirected graph of Boolean equivalences. However, because of noise the connected components become internally inconsistent e.g. two genes opposite to each other becomes part of the same connected component. In addition, the size of clusters became unusually big with almost everything in one cluster. To avoid such a situation, we need to break the component by removing the weak links. To identify the weakest links, we first computed a minimum spanning tree for the graph and computed Jaccard similarity coefficient for every edge in this tree. Ideally if two members are part of the same cluster they should share as many connections as possible. If they share less than half of their total individual connections (Jaccard similarity coefficient less than 0.5) the edges are dropped from further analysis. Thus, many weak equivalences were dropped using the above algorithm leaving the clusters internally consistent. We removed all edges that have Jaccard similarity coefficient less than 0.5 and built the connected components with the rest. The connected components were used to cluster the BIN which is converted to the nodes of the CBIN. The distribution of cluster sizes was plotted in a log-log scale to observe the characteristic of the Boolean network. The clusters sizes were distributed along a straight line in a log-log plot suggesting scale-free properties. The choice of the threshold on the Jaccard similarity coefficient play an important role in determining the size and the number of clusters as well as whether they are internally consistent. We found that a threshold of 0.5 gave us reasonable number of clusters and followed a scale-free distribution in the cluster sizes. A bigger threshold such as 0.7 to 0.9 will be very aggressive and reduce the cluster sizes (almost all edges will be dropped). A smaller number such as 0.4 will tend to make bigger cluster with unusual distribution of cluster sizes. A new graph was built that connected the individual clusters to each other using Boolean relationships. Link between two clusters (A, B) was established by using the top representative node from A that was connected to most of the member of A and sampling 6 nodes from cluster B and identifying the overwhelming majority of BIRs between the nodes from each cluster.

A CBIN was created using the selected Peters-2017 GSE83687 and Arijs-2018 GSE73661 datasets. Each cluster was associated with healthy or disease samples based on where these gene clusters were highly expressed. The edges between the clusters represented the Boolean relationships that are color-coded as follows: orange for low => high, dark blue for low => low, green for high => high, red for high => low, light blue for equivalent and black for opposite.

#### Generation of IBD, UC, and CD maps

IBD map is derived from the CBIN of the Peters-2017 GSE83687 and Arijs-2018 GSE73661 datasets by focusing on the largest clusters and their connections. A subset of the CBIN (**Supplemental Figure 1B**) is constructed by following the top 10 largest clusters and a Boolean path sequence of for high => high, high => low, and low => low (dark blue). Machine learning is performed on this network to identify Boolean path that can distinguish normal vs IBD samples. A Boolean path is converted to a path score as mentioned above using a linear combination of normalized gene expression values. The strength of classification of healthy and IBD samples using this score is computed by the ROC-AUC measurement. We performed a multivariate regression to identify the best Boolean path that predicts normal vs IBD samples in the cohort GSE6731 (4 N, 5 UC, 7 CD). Path #1-2-3 emerged as the winner. UC map (**Supplemental Figure 4**) and CD map (**Supplemental Figure 5**) are created by restricting the Peters-2017 GSE83687 dataset to UC only and CD only samples before constructing the CBIN respectively. Arijs-2018 GSE73661 dataset is not used for the UC and the CD maps.

#### Generation of heat maps and drug targets score

A composite score is computed as follows when many genes are considered drug targets which includes a summary of their gene expression values. To compute the composite score, gene expression values were normalized according to a modified Z-score approach centered around *StepMiner* threshold (formula = (expr - SThr)/3*stddev). The samples were ordered according to average of the normalized gene expression values in the given gene list. The heatmap use red colors for the high values, white colors for the intermediate values and blue colors for low values. Gene names for few selected genes are highlighted on the left to show their expression patterns. Drug targets score is computed as a linear combination of the normalized gene expression values (the modified Z-score as described above). Samples are ordered using the drug targets score and the strength of the association between gene expression and disease annotation is computed using ROC-AUC measurement. A barplot is used to visualize the sample ordering with different color codes for the disease annotation. Additionally, a set of violin plots is used just below the barplot to demonstrate the distribution of the drug target score across different disease annotations.

#### Identification of Epithelial-Mesenchymal and Inflammation-Fibrosis continuum

Top genes involved with Epithelial-Mesenchymal processes and inflammation-fibrosis processed are chosen from literature review, and used earlier (1). Given a list of genes *BoNE* computes a subgraph of the CBIN graph by identifying clusters that include one or more genes from this list. BoNE then search for a path in this subgraph as mentioned before with the original CBIN graph. The path identified is used to draw a model of the gene expression timeline. The continuum is identified by computing a score based on the path as described before.

#### GeneSet Enrichment Analysis (GSEA)

GeneSet Enrichment Analysis (GSEA) was performed using python gseapy 0.10.2 package. Difference in average expression values of two groups is used to compute gene rank file. GSEA pre-ranked analysis is performed on the precomputed rank file to check the significance of geneset enrichment score and generate the enrichment plot. GSEA computes four key statistics for the gene set enrichment analysis report: Enrichment Score (ES), Normalized Enrichment Score (NES), False Discovery Rate (FDR), Nominal P Value.

##### Measurement of classification strength or prediction accuracy

Receiver operating characteristic (ROC) curves were computed by simulating a score based on the ordering of samples that illustrates the diagnostic ability of binary classifier system as its discrimination threshold is varied along the sample order. The ROC curves were created by plotting the true positive rate (TPR) against the false positive rate (FPR) at various threshold settings. The area under the curve (often referred to as simply the AUC) is equal to the probability that a classifier will rank a randomly chosen IBD samples higher than a randomly chosen healthy samples. In addition to ROC AUC, other classification metrics such as accuracy ((TP + TN)/N; TP: True Positive; TN: True Negative; N: Total Number), precision (TP/(TP+FP); FP: False Positive), recall (TP/(TP+FN); FN: False Negative) and f1 (2 * (precision * recall)/(precision + recall)) scores were computed. Precision score represents how many selected items are relevant and recall score represents how many relevant items are selected. Fisher exact test is used to examine the significance of the association (contingency) between two different classification systems (one of them can be ground truth as a reference).

##### Statistical Analyses

All statistical tests were performed using R version 3.2.3 (2015-12-10). Standard t-tests were performed using python scipy.stats.ttest_ind package (version 0.19.0) with Welch's Two Sample t-test (unpaired, unequal variance (equal_var=False), and unequal sample size) parameters. Multiple hypothesis correction were performed by adjusting *p* values with statsmodels.stats.multitest.multipletests (fdr_bh: Benjamini/Hochberg principles). The results were independently validated with R statistical software (R version 3.6.1; 2019-07-05). Pathway analysis of gene lists were carried out via the Reactome database and algorithm(17). Reactome identifies signaling and metabolic molecules and organizes their relations into biological pathways and processes. Kaplan-Meier analysis is performed using lifelines python package version 0.22.8. Violin, Swarm and Bubble plots are created using python seaborn package version 0.10.1.

##### Code Availability

The codes are publicly available at the following links: https://github.com/sahoo00/BoNE; https://github.com/sahoo00/Hegemon

***Experimental methods***

**Reagents**

All reagents were purchased from Sigma-Aldrich (St. Louis, MO), unless otherwise indicated. Goat anti-rabbit and goat anti-mouse Alexa Fluor 680 and IRDye 800 F(ab')2 were purchased from LI-COR Biosciences (Lincoln, NE). Pioglitazone was purchased from Selleck Chemicals (Houston, TX). GW7647, GW6471 and GW9662 were purchased from Tocris Biosciences (Bristol, UK). PAR5359 was synthesized at Dr. Yang's lab, Department of Chemistry and Biochemistry, University of California San Diego.

**Synthesis of PAR5359**

*ethyl (S)-2-ethoxy-3-(4-(2-hydroxyethoxy) phenyl)propanoate* **(compound 1)**

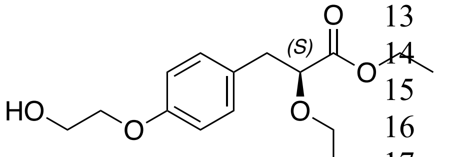


Ethylene carbonate (663 mg, 5.04 mmol, 3 equiv.) was added to the solution of ethyl (S)-2-ethoxy-3-(4 hydroxyphenyl) propanoate (400 mg, 1.68 mmol) and potassium carbonate ($K_2CO_3$) (695 mg, 5.04 mmol, 3 equiv.) in dry dimethylformamide (DMF) (5mL). The reaction was stirred at 80ºC overnight (16 hours). The reaction flask was then diluted with ~50 mL of ethyl acetate (EtOAc), and the solids were removed with filtration through celite. Water (~30 mL) was added, and the solution mixture was extracted twice with EtOAc (~50 mL x 2), the combined organics were washed with brine (~50 mL), dried over $MgSO_4$, filtered, and concentrated *in vacuo*. The product was purified via $SiO_2$ column chromatography (using a gradient of 20% to 30% to 50% EtOAc in hexanes as eluent) to give the title compound **1** as a clear oil (335 mg, **70% yield**). 1H NMR (300 MHz, $CDCl_3$) δ (ppm) = 7.16 (d, 2H), 6.84 (d, 2H), 4.16 (q, 2H), 4.04 (t, 2H), 3.98-3.91 (m, 3H), 3.64-3.54 (m, 1H), 3.38-3.28 (m, 1H), 2.95 (d, 2H), 2.23 (s, 1H), 1.21 (t, 3H), 1.15 (t, 3H).

*ethyl (S)-2-ethoxy-3-(4-(2-((methylsulfonyl)oxy)ethoxy)phenyl)propanoate* **(compound 2)**

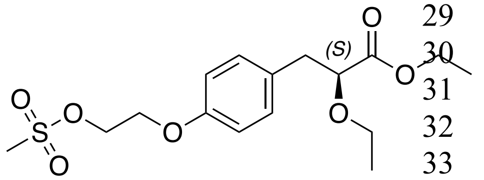


Methanesulfonyl chloride (MsCl, 237 mg, 0.16 mL, 2.02 mmol, 1.7 equiv.) was added dropwise to an ice-cold solution of compound **1** (335 mg, 1.19 mmol) and triethylamine (TEA) (240 mg, 0.331 mL, 2.38 mmol, 2 equiv.) in dry dichloromethane (DCM) (7mL). The reaction was stirred at room temperature for 2.5 hours, then diluted with ~50 mL of 1M HCl aq. solution. The aqueous layer was then extracted with DCM (50mL x 2), the combined organic layers were washed with sequence of ~50 mL of saturated $NaHCO_3$ aq. solution, ~50 mL of water, and ~50mL of brine. The organic layer was dried over $MgSO_4$, filtered, and concentrated *in vacuo*, to give the title compound **2** as a brown oil, with no further purification (412 mg, **96% yield**). 1H NMR (300 MHz, $CDCl_3$) δ ppm = 7.19 (d, 2H), 6.83 (d, 2H), 4.57 (t, 2H), 4.22 (t, 2H), 4.16 (t, 2H), 3.97 (q, 1H), 3.66-3.55 (m, 1H) 3.40-3.30 (m, 1H), 3.09 (s, 3H), 2.97 (d, 2H), 1.24 (t, 3H), 1.16 (t, 3H)

*ethyl (S)-3-(4-(2-(4-(4-chlorophenyl)-3,6-dihydropyridin-1(2H)-yl)ethoxy)phenyl)-2-ethoxypropanoate* (**compound 3**)

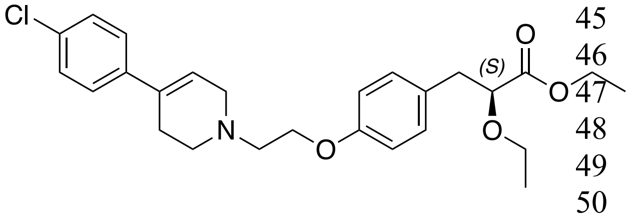

*4-(4-chlorophenyl)-1,2,3,6tetrahydropyridine* (314 mg, 1.37 mmol, 1.2 equiv.), sodium iodide (NaI) (34 mg, 0.23 mmol, 0.2 equiv.), and potassium carbonate ($K_2CO_3$) (471 mg, 3.42 mmol, 3 equiv.) was added to the solution of compound **2** (412 mg, 1.14 mmol) in dry DMF (6 mL). The reaction was stirred at 60ºC overnight (16 hours). The reaction flask was then diluted with ~50 mL of EtOAc, and the solids were removed by filtration through celite. Water (~30 mL) was added, and the solution mixture was extracted three times with EtOAc (~50 mL x 3), the combined organics were washed with brine (~50 mL), dried over $MgSO_4$, filtered, and concentrated *in vacuo*. The product was purified via $SiO_2$ column chromatography (using a gradient of 20% to 30% to 40% EtOAc in hexanes as eluent) to give the title compound **3** as a clear oil (189 mg, **36% yield**). 1H NMR (300 MHz, $CDCl_3$) δ ppm = 7.33 (d, 2H), 7.28 (d, 2H), 7.17 (d, 2H), 6.86 (d, 2H), 6.06 (s, br, 1H), 4.20-4.08 (m, 3H), 3.97 (t, 1H), 3.65-3.55 (m, 1H), 3.40-3.32 (m, 1H), 3.31 (d, 2H), 2.94 (t, 4H), 2.86 (t, 2H), 2.57 (s, br, 2H), 1.23 (t, 3H), 1.16 (t, 3H).

*(S)-3-(4-(2-(4-(4-chlorophenyl)-3,6-dihydropyridin-1(2H)-yl)ethoxy)phenyl)-2-ethoxypropanoic acid* (**PAR5359**)

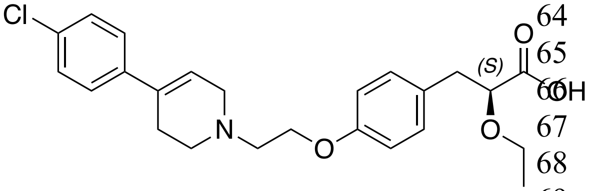

Lithium hydroxide monohydrate (26 mg, 0.624 mmol, 2 equiv.) was added to the solution of compound **3** (163 mg, 0.312 mmol) in tetrahydrofuran (THF) (6 mL) and water (1.5mL). The reaction was stirred at room temperature for 4 hours and was then quenched by addition of ~1 mL 1M HCl aq. solution. The reaction flask was then evaporated to dryness *in vacuo*. The resultant solids were purified via $SiO_2$ column chromatography (using a gradient of 4% to 6% to 10% MeOH in DCM as eluent) to give the title compound **PAR5359** as a white solid (109 mg, **71% yield**) 1H NMR (500 MHz, CD3OD-d4) δ ppm = 7.25-7.23 (d, 2H), 7.13-7.11 (d, 2H), 6.98-6.97 (d, 2H), 6.70-6.68 (d, 2H), 5.94 (t, J= 2.2 Hz, 1H), 4.15-4.13 (m, 2H), 3.74-3.73 (m, 2H), 3.60-3.59 (m, 2H), 3.38-3.32 (m, 5H), 3.05-2.98 (m, 1H), 2.74-2.71 (m, 1H), 2.63-2.55 (m, 3H), 0.84 (t, J= 7.0 Hz, 3H); 13C NMR (126 MHz, CD3OD-d4) δ ppm = 181.1, 158.7, 139.6, 136.4, 135.7, 134.2, 132.5, 132.4, 130.6, 130.5, 128.7, 128.6, 119.0, 118.9, 116.2, 116.2, 84.6, 67.4, 64.7, 57.1, 53.3, 51.7, 40.6, 26.5, 16.3; ESI-MS: 430.2 $[M+H]^+$.

**Bacteria and bacterial culture**

For bacterial culture adherent Invasive *Escherichia coli* strain LF82 (AIEC-LF82), *Citrobacter rodentium* and *Salmonella enetrica serovar typhimurium*, a single colony was inoculated into LB broth and grown for 6-8 h on shaking incubator, followed by overnight culture under oxygen-limiting conditions, but without shaking, to maintain their pathogenicity as done previously(18-20). Cells were infected with bacteria with indicated MOI in figure legends.

***C. rodentium* induced infectious colitis and *in vivo* treatments**

*C. rodentium* were grown overnight in LB broth with shaking at 37 ºC and mice were orally challenged with 5 x $10^8$ CFU in 0.1 ml of PBS as described (21, 22). To determine viable bacterial numbers in faeces, fecal pellets were collected from individual mice, homogenized in cold PBS, serially diluted and plated on MacConkey agar plates. Number of CFU was determined after overnight incubation at 37 ºC. Colon samples were collected to assess histology and levels of mRNA (by qPCR). For treatment study, PPARα agonist (GW7647, 20 μg/kg body weight/day), PPARγ agonist (Pioglitazone, 20μg/kg body weight/day), PPARα/γ dual agonist (PAR5359, 1 mg/kg body weight/day) were administered via intraperitoneal route in 200 μl total volume (DMSO less than 4%).

**DSS-induced colitis and *in vivo* treatments**

For DSS-colitis experiments, 7-wk old C57BL/6 mice were obtained from Jackson Laboratories (Bay Harbor, ME). All animals were housed and euthanized according to University of California San Diego Institutional Animal Care and Use Committee (IACUC) policies and guidelines. Colitis was induced by oral administration of 2.5% dextran sulfate sodium (DSS, w/v) (MP Biomedicals, MW 36–50 kDa) in drinking water for five days as described (23, 24). For treatment study, PAR5359 (1 mg/kg/day) was administered via intrarectal/intraperitoneal route in 50 μl total volume (DMSO less than 4%). Post-injection, mice were hung upside-down for 30 sec to ensure injection solution was retained in colon. Mice were sacrificed on the 9th day, and colon length was assessed. Colon samples were collected for assessing the levels of mRNA (by qPCR). Water levels were monitored to determine the volume of water consumed by all groups. Each animal was monitored for animal weight loss, stool consistency, and fecal blood and these parameters were used to calculate an average Disease Activity Index (DAI) as described previously (25). Colon histology was assessed in samples stained with hematoxylin using standard protocols.

**Thioglycolate-elicited murine peritoneal macrophages generations**

Thioglycolate-elicited murine peritoneal macrophages (TGPMs) were isolated from 8- to 12-week-old C57BL/6 mice and cultured as described previously(26). Peritoneal cells were collected from peritoneal lavage with ice cold RPMI (10 ml per mouse) 4 days after intraperitoneal injection of 3 ml of aged, sterile 3% thioglycolate broth (BD Difco, USA). Cells were filtered with 70 μ filter, centrifuged and resuspended in RPMI-1640 containing 10 % FBS and 1% penicillin/streptomycin. TGPMs were plated with required cell density and the media was changed after 4 h to remove non adherent cells. Depending on the experiment, TGPMs were seeded in 6-well, 12-well, or 24-well plates with appropriate and consistent cell densities. TGPMs were allowed to adjust to overnight culture before the addition of stimuli: LPS (10-100 ng/ml) in presence or absence of PPAR agonists and antagonist as describe in figure legends.

**Measurement of reactive oxygen species**

To assess whether PPAR agonists modulates bacteria (LF82 and SL)-induced ROS in peritoneal macrophages were (50,0000 cells/96 well) were treated with *AIEC*-LF82/SL in presence or absence of PPAR agonists/inhibitors. The redox-sensitive, cell-permeable dihydroethidium (hydroethidine or DHE) was used to detect the cellular production of ROS as described in assay kit (ROS, Detection Cell-Based Assay Kit, Cayman Chemical) and plate was read using fluorescence microplate reader (ex 500 – 530 nm/em 590 – 620 nm).

**Gentamicin Protection Assay.**

Quantification of viable intracellular bacteria was done by using the gentamicin protection assay as described previously (18). Briefly, Peritoneal macrophage TGPMs, $2 \times 10^5$ cells per well were seeded into 24-well culture dishes overnight before infection at an MOI of 10 for 1 h in antibiotic-free RPMI media containing 10% FBS in a 37 °C CO2 incubator. Cells were then washed and incubated with gentamicin (200 µg/ml) for 90 min to kill extracellular bacteria. Further, cells were washed and incubated with antibiotic-free RPMI media containing 10% FBS for 1-24 h and subsequently lysed in 1% Triton-X 100, lysates were serially diluted and plated on LB agar plates. Total CFU were counted after overnight incubation at 37 °C. To test effect of PPAR agonists, cells were pre-treated for 30 min with PPAR agonists.

**RNA extraction and Quantitative PCR**

Total RNA was isolated using TRIzol reagent (Life Technologies) and Quick-RNA MiniPrep Kit (Zymo Research, USA). RNA was converted into cDNA using the qScript™ cDNA SuperMix (Quantabio). Quantitative RT-PCR (qPCR) was carried out in 384-well plate using PowerUp™ SYBR™ green master mix (Applied Biosystems, USA) and performed on the SteepOnePlus Quantitative platform (Life Technologies, USA). The cycle threshold (Ct) of target genes was normalized to 18S rRNA gene and the fold change in the mRNA expression was determined using the 2^-ΔΔCt method. Primers used in qPCR reactions were designed using PrimerQuest tool software (Integrated DNA technology, USA) and NCBI Primer Blast software (Table 1).

**RNA-seq library preparation**

Sequencing libraries were generated using the Illumina TruSeq Stranded Total RNA Library Prep Gold with TruSeq Unique Dual Indexes (Illumina, San Diego, CA) as previously described (27). Libraries were amplified and sequenced on an Illumina NovaSeq 6000 by the Institute of Genomic Medicine (IGM) at the University of California San Diego.

**Cytokine Assays**

Cyokines including TNFa, IL6, IL1b and IL10 were measured in cell supernant using ELISA MAX Deluxe kits from Biolegend.

**Statistics**

Experimental values are presented as the means of replicate experiments ±SEM. Statistical analyses were performed using GraphPad Prism software version 8.0 (GraphPad Software). Differences between two groups were evaluated using Student's t-test (parametric) or Mann-Whitney U-test (non-parametric). To compare more than three groups, one-way analysis of variance followed by Tukey's post-hoc test was used. Differences at $P < 0.05$ were considered significant. Please see Supporting Information for details regarding Boolean data analysis.

**Study approval**

***Human subjects.*** Blood samples were obtained from either healthy volunteers or from IBD patients undergoing colonoscopies a part of their routine care and follow-up at UC San Diego's

Inflammatory Bowel Disease (IBD) Center. Patients were recruited and consented using a study proposal approved by the Institutional Review Board of the University of California, San Diego. Isolation of blood monocytes was carried out using an approved human research protocol (IRB# 160246) that covers human subject research at the UC San Diego HUMANOID Center of Research Excellence (CoRE). The clinical phenotype and information were curated based on histopathology reports from Clinical Pathology and Chart check, followed by consultation with a specialist at UC San Diego's IBD Center.

***Animals.*** All animal studies were approved by the University of California, San Diego Institutional Animal Care and Use Committee (IACUC). Adult C57BL/6 mice were acquired from Jackson Laboratories. All animals were maintained in an institutional animal care. Provided with standard light–dark cycle, fed with standard laboratory chow and clean drinking water.

#### KEY RESOURCE TABLE

| REAGENT or RESOURCE | SOURCE | IDENTIFIER |
|---|---|---|
| ***Biological Samples and Cell Lines*** | | |
| *PPARα reporter assay system* | Indigo Biosciences | IB00111-32 |
| *PPARγ reporter assay system* | Indigo Biosciences | IB00101-32 |
| *E. coli LF82 (AIEC-LF82)* | Prof. Arlette Darfeuille-Michaud | (28) |
| *Salmonella enteric serovar typhimurium strain SL1344* | ATCC | 700720 |
| ***Chemicals and Reagents*** | | |
| *PAR5359* | This study (see *Methods*) | |
| *GW7647 (PPARα agonist)* | Tocris bioscience | 1677 |
| *Pioglitazone (PPARγ agonist)* | Selleck Chemicals | S2590 |
| *GW 6471 (PPARα antagonist)* | Tocris bioscience | 4618 |
| *GW 9662 (PPARγ antagonist)* | Tocris bioscience | 1508 |
| *Lipopolysaccharide (E. coli O111:B4)* | Sigma-Aldrich | L4391 |
| *PowerUpä SYBRä Green Master Mix* | Applied Biosciences | A25741 |
| *qScriptä cDNA SuperMix* | QuantaBio | 101414 |
| *Direct-zol RNA Miniprep Kit* | Zymo Research | R1051 |
| *TRIzolä Reagent* | Invitrogen | 15596018 |
| *ELISA MAXä Deluxe Set Mouse IL-6* | BioLegend | 431304 |
| *ELISA MAXä Deluxe Set Mouse IL-1b* | BioLegend | 432604 |
| *ELISA MAXä Deluxe Set Mouse IL-10* | BioLegend | 431414 |
| *ELISA MAXä Deluxe Set Mouse TNF-α* | BioLegend | |

| | | |
|---|---|---|
| *Dextran Sulfate Sodium Salt (Colitis Grade)* | MP Biomedicals, LLC | 160110 |
| *Hemoccult II* | Beckman Coulter | 61130 |
| *Zinc Formalin Fixative* | Sigma-Aldrich | Z2902 |
| *ROS Detection Cell-Based Assay Kit (DHE)* | Cayman Chemical | 601290 |
| ***Primers*** | | |
| *Species/Targets* | Forward primer (5'→ 3') | Reverse primer (3'→ 5') |
| *Mouse IL-6 qPCR primers* | TGGAGTCACAGAAGGAGTGGCTAAG | TCTGACCACAGTGAGGAATGTCCAC |
| *Mouse IL-1b qPCR primers* | GCCTTGGGCCTCAAAGGAAAGAATC | GGAAGACACAGATTCCATGGTGAAG |
| *Mouse TNFa qPCR primers* | ATAGCTCCCAGAAAAGCAAGC | CACCCCGAAGTTCAGTAGACA |
| *Mouse IL-10 qPCR primers* | CCCTGGGTGAGAAGCTGAAG | CACTGCCTTGCTCTTATTTTCACA |
| *Mouse 18S qPCR primers* | GTAACCCGTTGAACCCCATT | CCATCCAATCGGTAGTAGCG |
| *Human PPARA* | CATTACGGAGTCCACGCGT | ACCAGCTTGAGTCGAATCGTT |
| *Human PPARG* | GAGAAGGAGAAGCTGTTGGC | ATGGCCACCTCTTTGCTCT |
| *Human PPARGC1A* | GCTACGAGGAATATCAGCACGA | ACACGGCGCTCTTCAATTG |
| ***Software*** | | |
| *Prism* | GraphPad | https://www.graphpad.com/scientific-software/prism/ |
| *Illustrator* | Adobe | https://www.adobe.com/products/illustrator.html |
| *ImageStudio Lite* | LI-COR | https://www.licor.com/bio/image-studio-lite/ |

#### SUPPLEMENTARY FIGURES

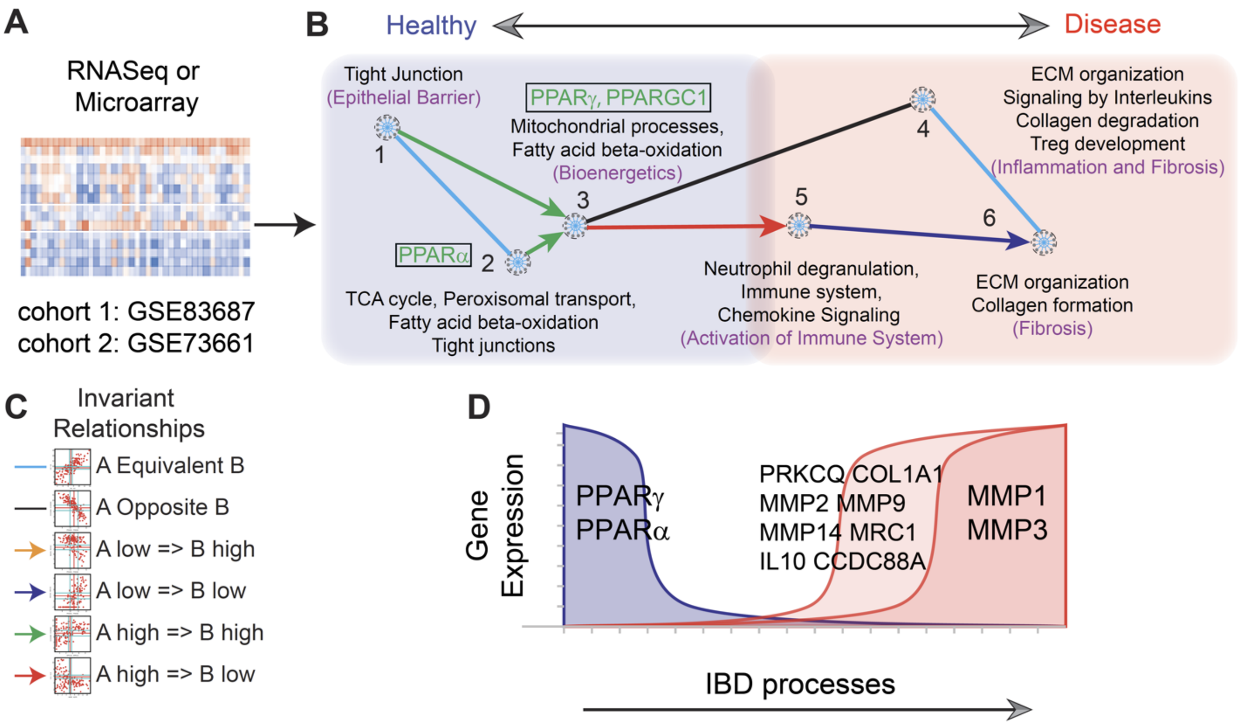


**Supplementary Figure 1: Boolean network map of continuum states in Inflammatory Bowel Disease (a.k.a, IBD-map) and the position of PPARA/G targets within the map.**
(**A**) Boolean network analysis was performed on IBD datasets (GSE83687 and GSE73661) to identify pathways and gene clusters during IBD progression. (**B**) Genes with similar expression profiles were organized into clusters, and relationships between clusters represented as color-coded edges connecting clusters. Reactome pathway of each cluster of IBD-map was performed to understand pathophysiological cellular processes that are enriched during IBD progression. PPARA is present within cluster #2 and PPARG is present within cluster #3. (**C**) Boolean networks contain six possible Boolean relationships between genes (invariant relationships). (**D**) Schematic showing the gene expression of selected genes within the normal to IBD disease progression.

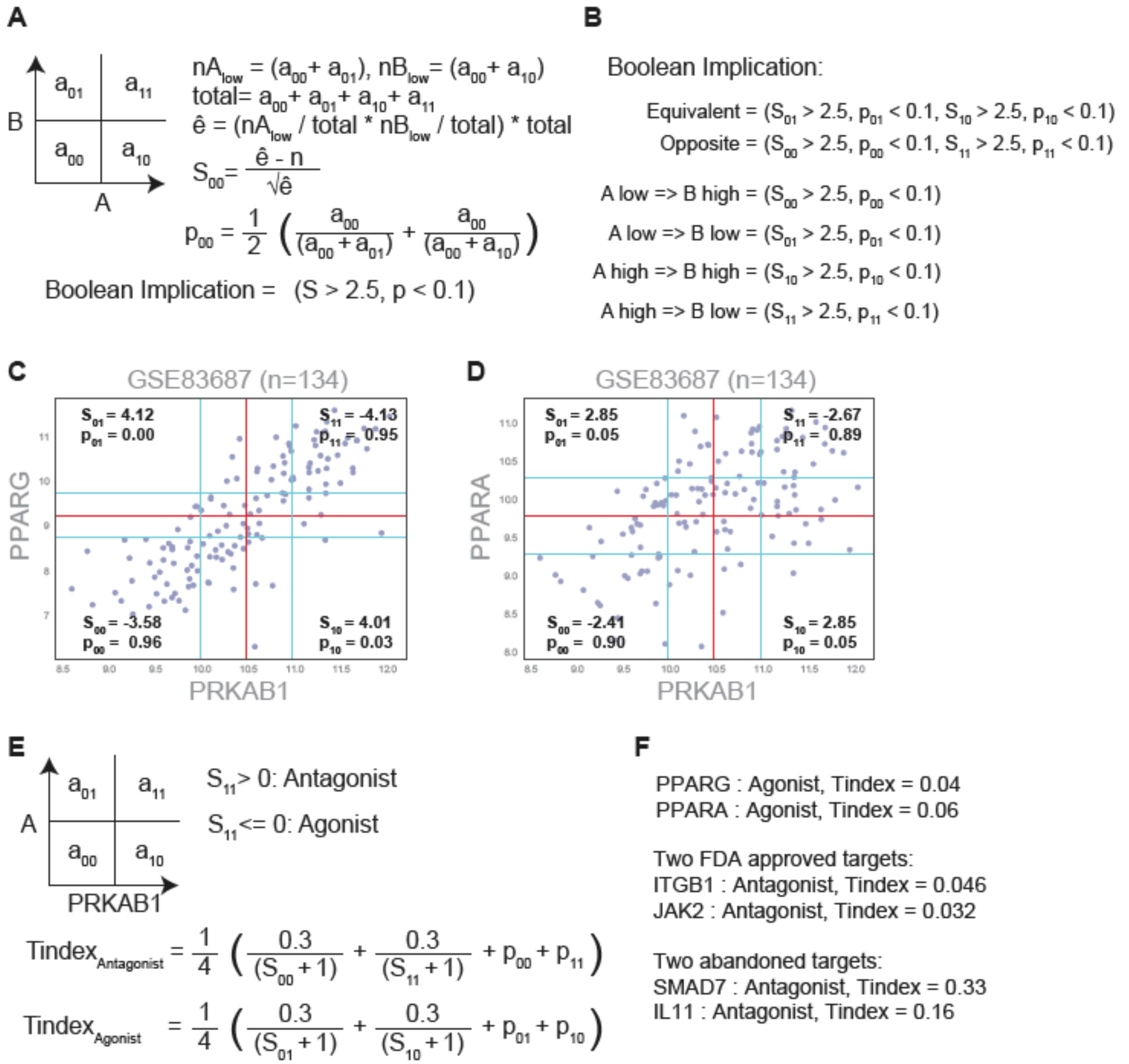


**Supplementary Figure 2: Computation of therapeutic index in a target report card.**
**(A)** BooleanNet statistic. Evaluating Boolean implication relationship between gene A and B. $a_{ij}$ is the number of samples in the respective quadrants. $nA/B_{low}$ is number of samples where A/B is low. $S_{00}$ = BooleanNet statistic and $p_{00}$ = error rate to test sparsity for the bottom left quadrant. $S > 2.5$ and $p < 0.1$ is used to test whether each quadrant is sparse. False Discovery Rate is computed by randomizing the data several times and computing the ratio of an average number of relationships found in randomized data to the original data. **(B)** Deriving Boolean implication relationships using BooleanNet statistic. **(C-D)** Scatterplots between PRKAB1, PPARG (C) and PPARA (D) in GSE83687 (n = 134) with the StepMiner thresholds (red) and noise margin (+/- 0.5, blue) in both X and Y-axes. BooleanNet statistic (S, p) is computed for each quadrant. **(E)** Since PRKAB1 is an Agonist, gene A is considered Antagonist if $S_{11} > 0$ (Top-right quadrant have fewer points than expected) and Agonist otherwise. The therapeutic index (Tindex) is computed separately for Antagonist and Agonist as shown below. **(F)** The therapeutic index (Tindex) value of PPARG, PPARA, two FDA approved targets (ITGB1, JAK2), two abandoned targets (SMAD7, IL11) is computed using the formula in panel E.

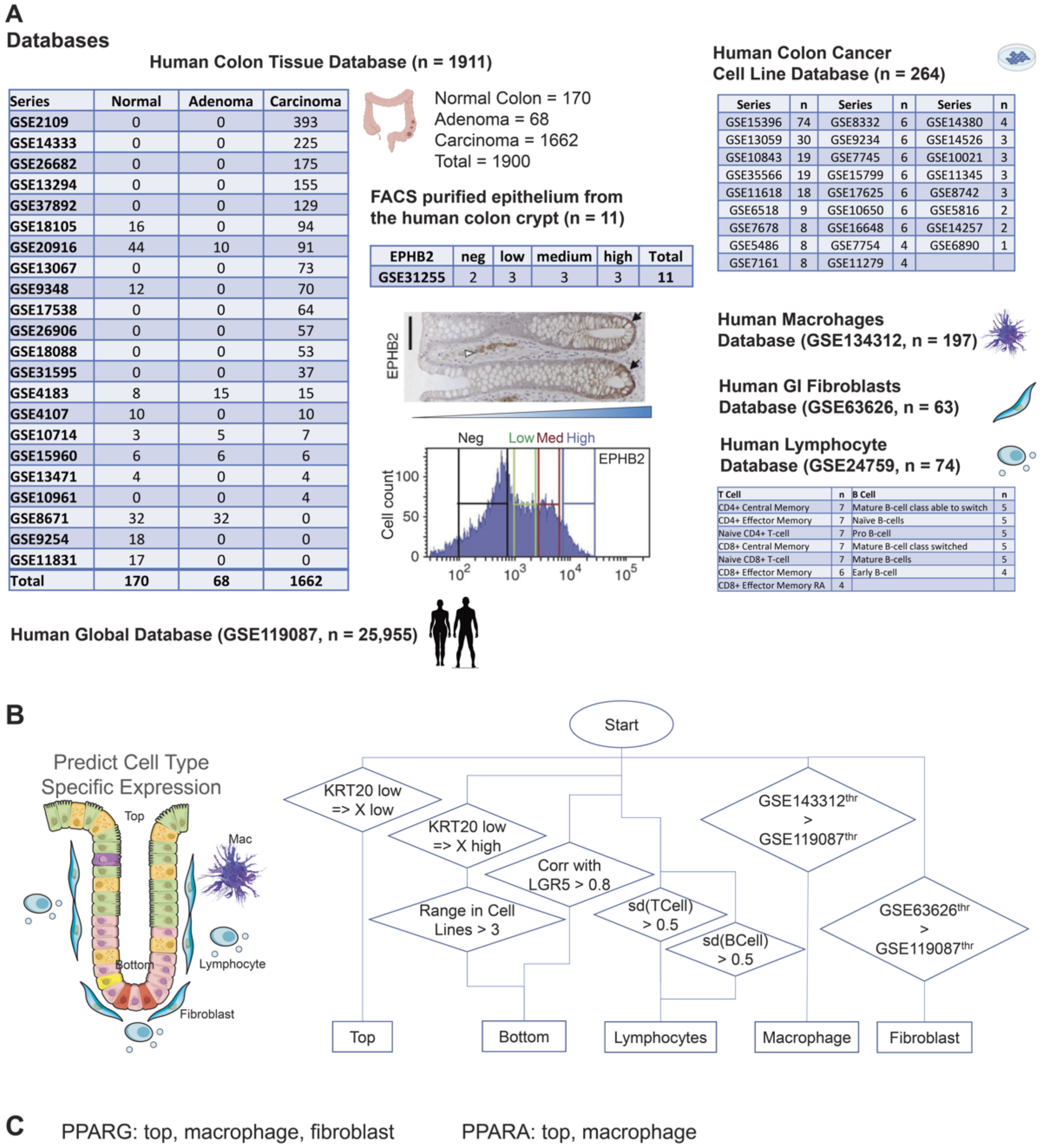


**Supplementary Figure 3: Prediction of cell type specific expression patterns.** **(A)** Gene expression databases used for predicting cell type specific expression patterns. Human colon tissue database (n = 1911) is derived from the "Human Colon Global Database after purging using EpCAM and Albumin" restricted to Human U133 Plus 2.0 Affymetrix platform as published previously (Dalerba, Sahoo et al. 2016, NEJM, PMID: 26789870) with additional 68 Adenoma samples and purified FACS samples of human colon crypts (GSE31255). A database of human colon cancer cell lines (n = 264) was prepared by pooling 26 independent datasets. Macrophage, and Fibroblast databases were prepared from GSE134312 and GSE63626, respectively. B cell and T cell samples from GSE24759 was used to prepare the human Lymphocyte database. Human Global Database (GSE119087, n = 25,955) is used to identify high and low expression patterns relative to other tissue types for each gene. **(B)** Flow chart for the prediction of cell type specific expression patterns. Boolean implication "KRT20 low => X low" in the human colon tissue database (n = 1911) is used to identify the top of the crypt specific expression patterns. Boolean implication

“KRT20 low => X high” in the human colon tissue database (n = 1911), dynamic range > 3 in human colon cancer cell lines (n = 264), and correlation with LGR5 > 0.8 in GSE31255 is used to identify bottom of the crypt specific expression patterns. Standard deviation > 0.5 in B cell and T cell samples from GSE24759 is used to predict if a gene is expressed in lymphocytes. Macrophage and Fibroblast specific expression pattern is predicted by comparing StepMiner thresholds of GSE134312 and GSE63626 with GSE119087. **(C)** Cell type specific expression patterns for PPARG and PPARA is computed using the flow chart in panel B.

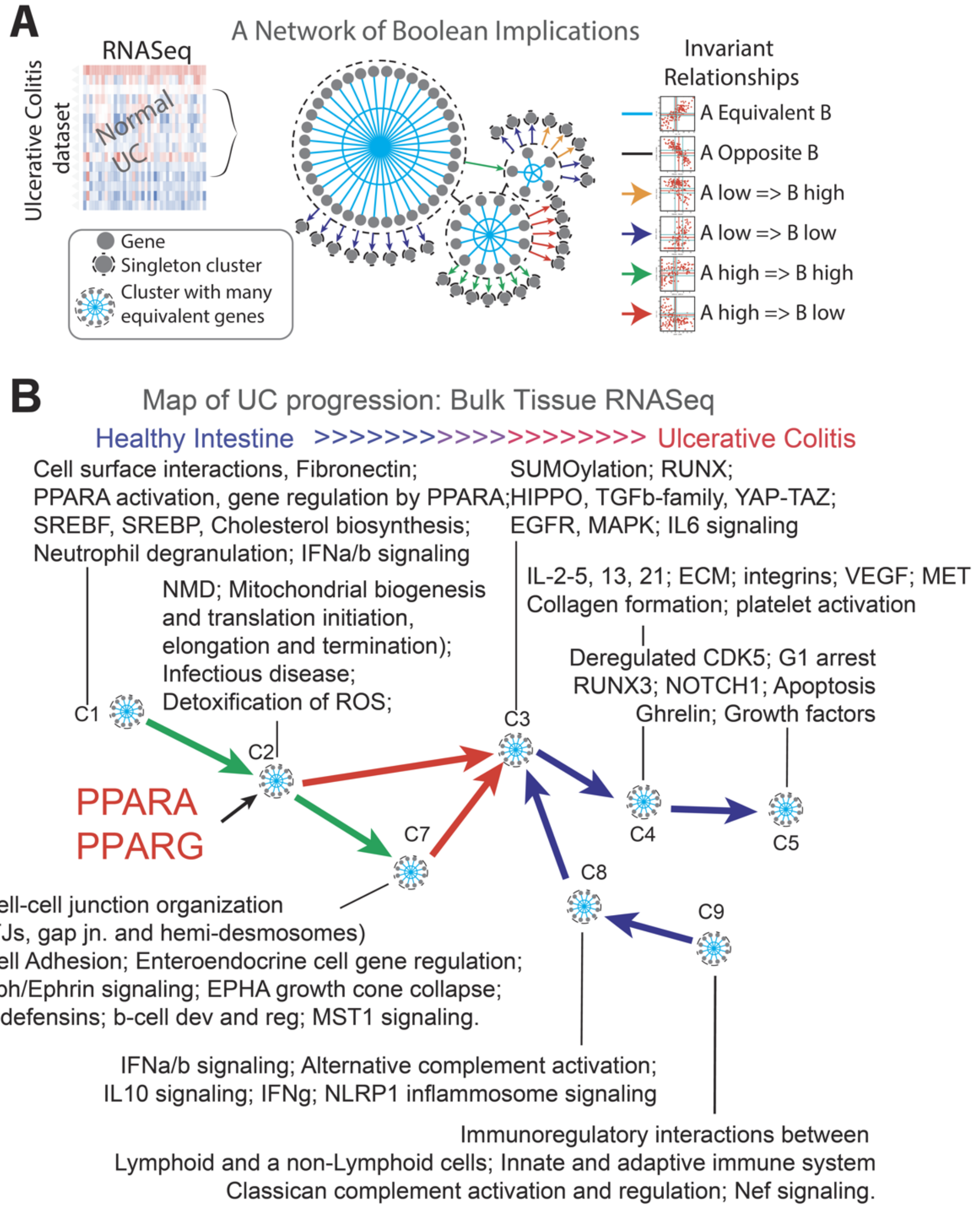


**Supplementary Figure 4: Boolean network map of continuum states in Ulcerative Colitis (a.k.a, UC-map) and the position of PPARA/G targets within the map.**
(A) Boolean network analysis was performed on Ulcerative colitis datasets to identify pathways and gene clusters during UC progression. Boolean networks contain six possible Boolean relationships between genes (invariant relationships top right) and genes with similar expression profiles were organized into clusters, and relationships between clusters represented as color-coded edges connecting clusters. (B) Schematic of reactome pathway of each cluster of UC-map were performed to understand pathophysiological cellular processes that are enriched during UC progression. PPARA and PPARG are both located in cluster #2. Their stimulation with dual agonists is expected to impact both continuum paths, C2→C3→C4→C5 and C2→C7→C3→C4→C5.

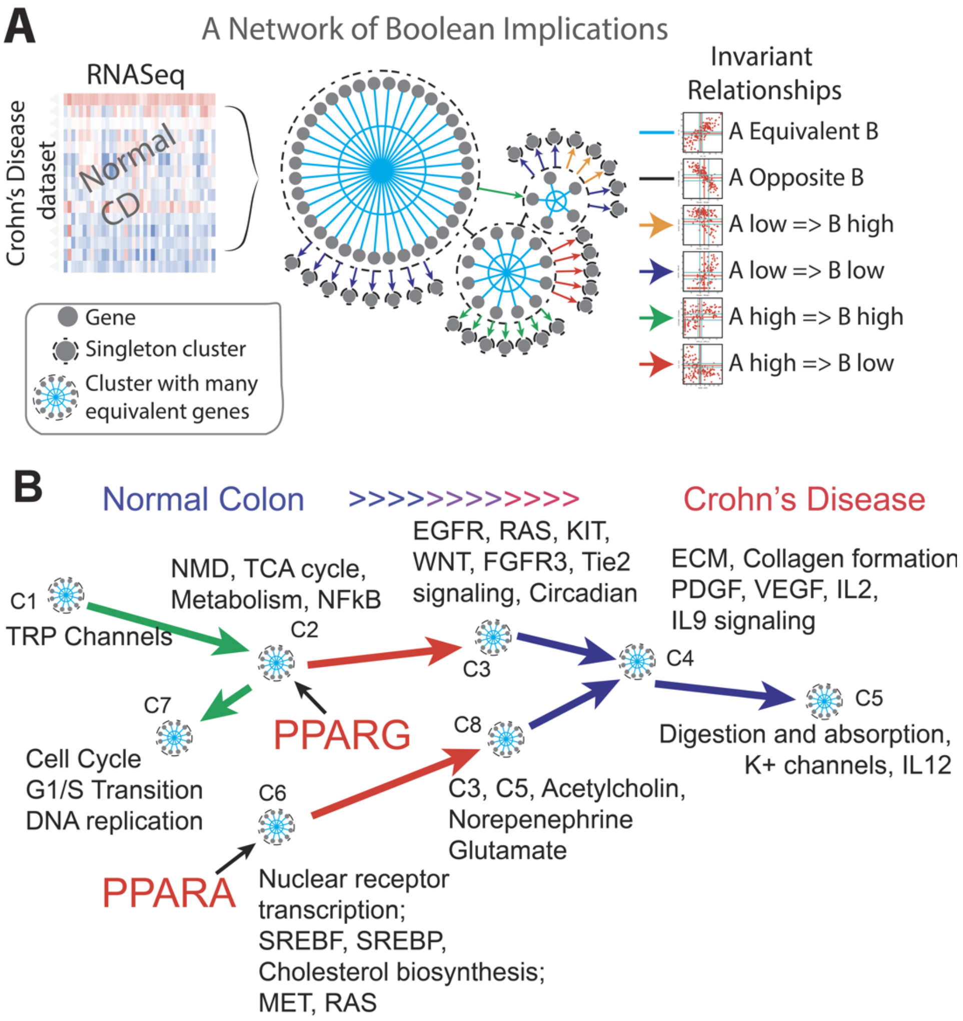


**Supplementary Figure 5: Boolean network map of Crohn's disease, CD-map.**
A) Boolean network analysis was performed on Crohn's colitis datasets to identify pathways and gene clusters during CD disease progression. Boolean networks contain six possible Boolean relationships between genes (invariant relationships top right) and genes with similar expression profiles were organized into clusters, and relationships between clusters represented as color-coded edges connecting clusters. (B) Schematic of reactome pathway of each cluster of CD-map were performed to understand pathophysiological cellular processes that are enriched during CD progression. PPARA is located within cluster #6 and PPARG is located within cluster #3. Their stimulation with dual agonists is expected to impact both continuum paths, C2→C3→C4→C5 and C6→C8→C4→C5.

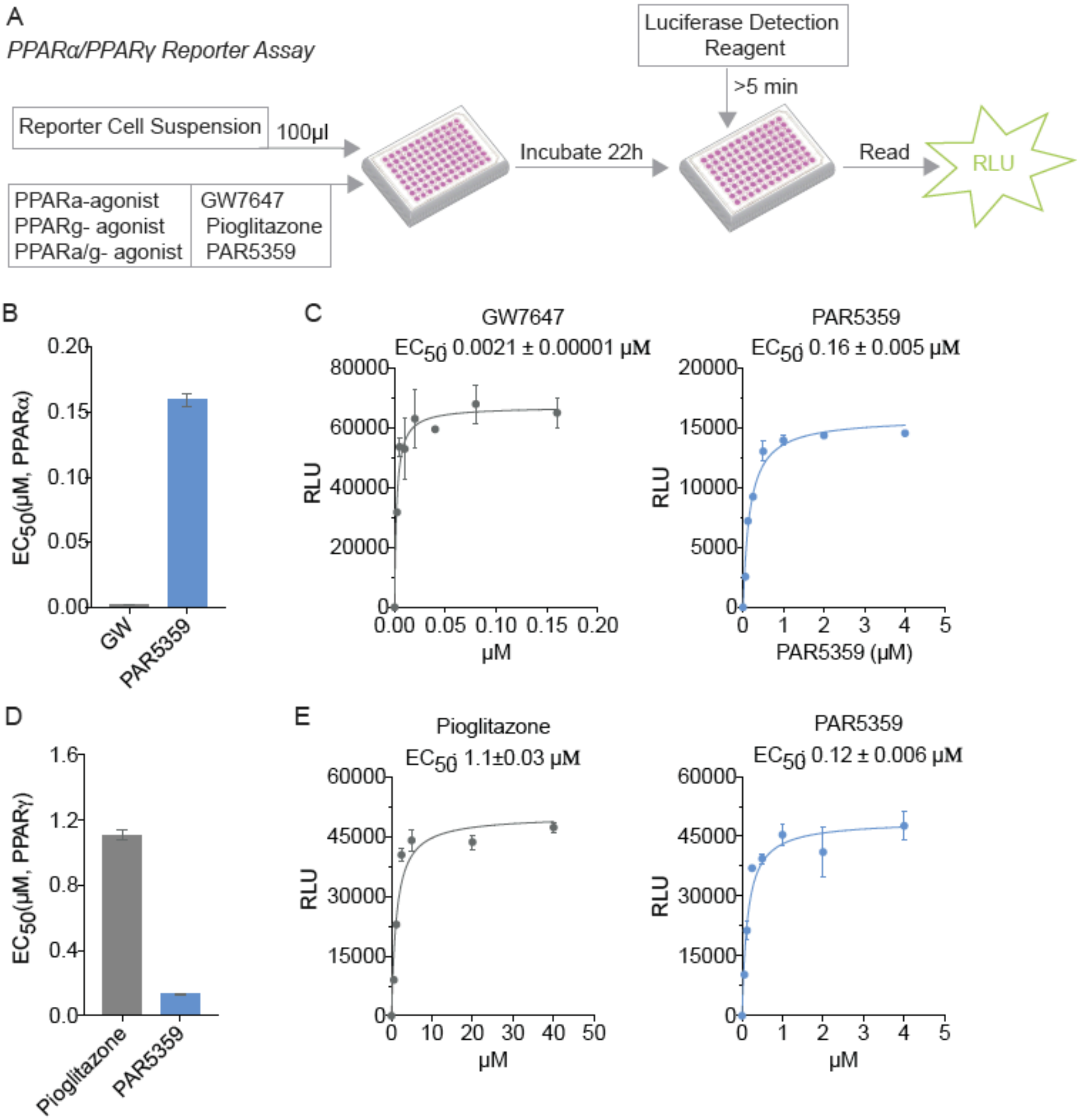


**Supplementary Figure 6**: **A comparative analysis of potencies of PPARα and PPARγ single and dual agonists.**

(A) Schematic displaying assay workflow. PPARα and PPARγ reporter cells were dispensed into respective wells of the assay plate and incubated with PPAR agonists (GW7647 and PAR5359 to test PPARα agonist activity and Pioglitazone and PAR5359 to test PPARγ agonist activity). Following 22 h incubation with agonists, treatment media are discarded, and Luciferase Detection Reagent was added (as indicated by manufacture's protocol, Indigo Biosciences). The intensity of light emission (in units of 'Relative Light Units'; RLU) from each assay well was quantified using a plate-reading luminometer. (B-C) Bar graph and line graph showing $EC_{50}$ for PPARα and (D-E) showing $EC_{50}$ for PPARγ.

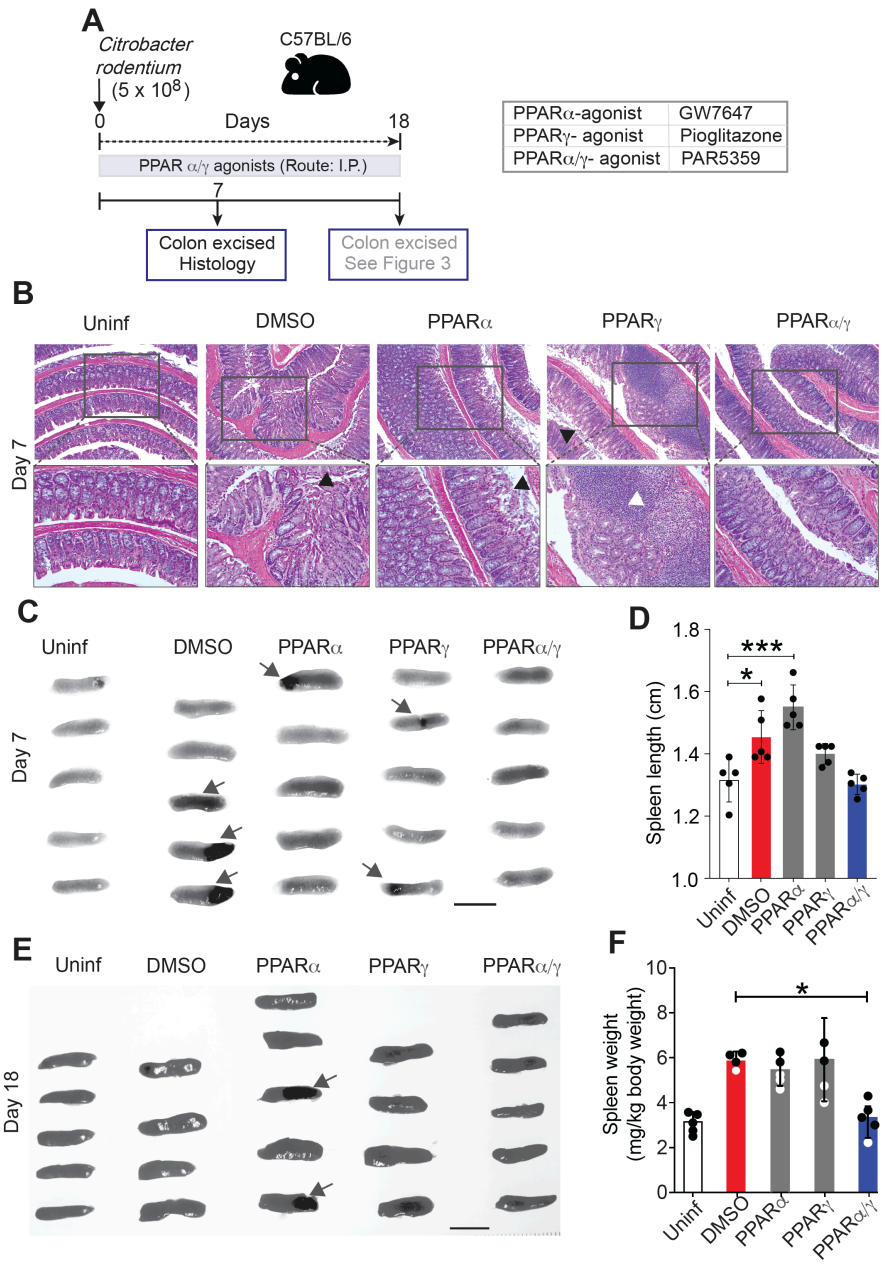


**Supplementary Figure 7**: **PPARα/γ dual agonists ameliorate *Citrobacter rodentium*-induced infectious colitis in mice (as determined on day #7, i.e., peak inflammation). (A)** Schematic summarizing the workflow for testing PPAR-targeted therapeutics in *C. rodentium*-induced colitis. Mice were gavaged with *C. rodentium* on day 0 and subsequently treated daily with PPAR agonists. Colons were excised on day 7 and analyzed by histology. **(B)** Images display representative fields from H&E-stained colon tissues. Mag = 100x (top) and 200x (bottom). White arrowheads point to immune cell infiltrates, whereas black arrowheads point to regions of extensive epithelial destructions and sloughing. **(C-D)** Images in panel C display spleens excised on day 7. Arrows point to black discoloration, likely from splenic infarcts. Scatter plots with bar graphs in panel D display the length of the spleens in C. (**E-F**) Images in panel E display spleens excised on day 18. Arrows point to black discoloration, likely from splenic infarcts. Scatter plots with bar graphs in panel F display the weight of the spleens in E. Statistics: All results are displayed as mean ± SEM. Significance was tested using two-way/one-way ANOVA followed by Tukey's test for multiple comparisons. Significance: *, $p < 0.05$Statistics: All results are displayed as mean ± SEM. Significance was tested using two-way/one-way ANOVA followed by Tukey's test for multiple comparisons. Significance: *, $p < 0.05$; ***, $p < 0.001$. See also Figure 4 for the Day 18 results in the *C. rodentium*-induced colitis model

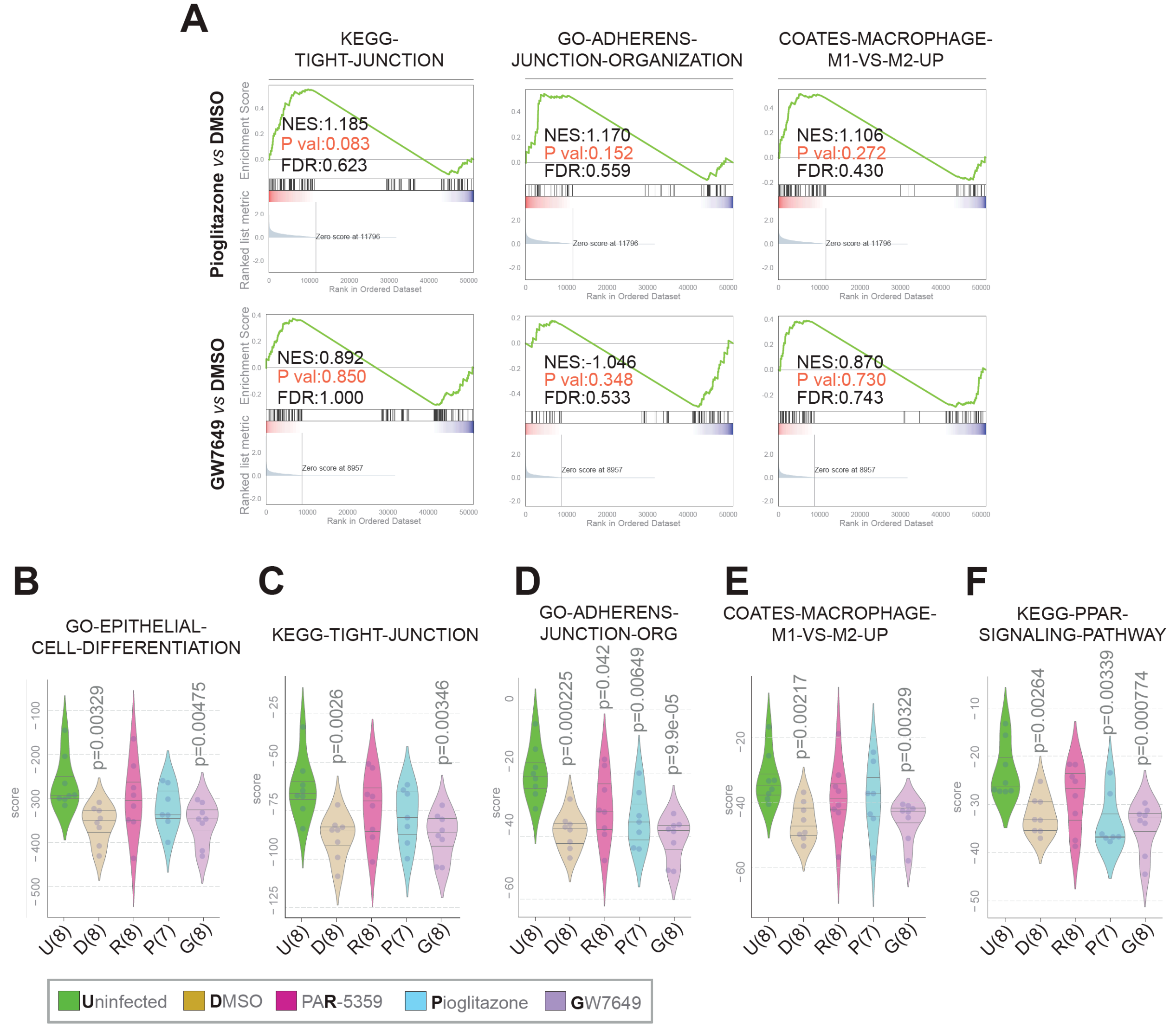


**Supplementary Figure 8**: **RNA seq analyses of *C. rodentium*-infected colons show that dual agonist PAR5359, but not single agonists GW7647 or Pioglitazone resists *Citrobacter rodentium*-induced gene expression changes.** (**A**) Pre-ranked GSEA based on pairwise differential expression analyses (Pioglitazone vs DMSO, *top*; GW7647 vs DMSO, *bottom*) are displayed as enrichment plots for epithelial tight (left) and adherens (middle) junction signatures and balanced macrophage processes (right). (**B-F**) Violin plots display the deviation of expression of gene sets that represent epithelial differentiation (B), epithelial tight and adherens junctions (C-D), macrophage processes (E) and PPAR signaling (F).

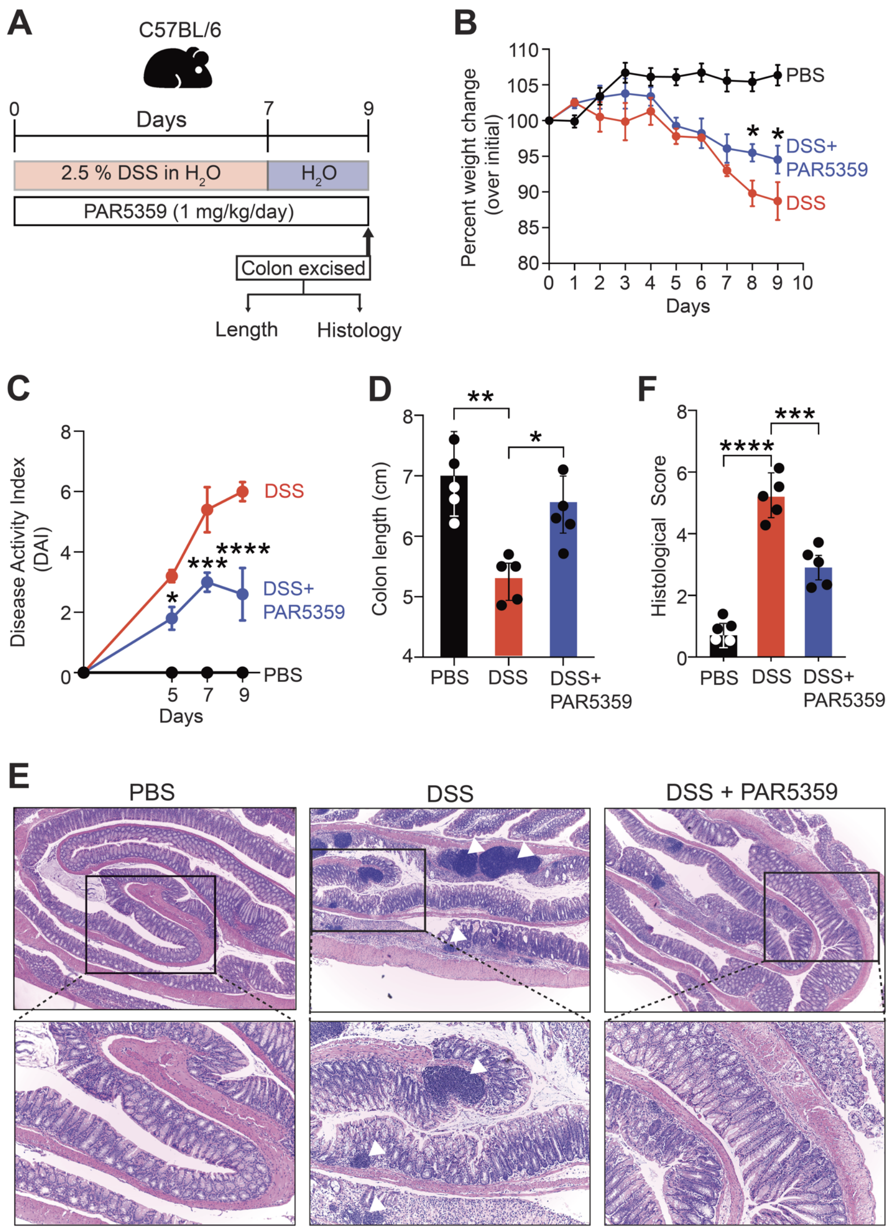


**Supplementary Figure 9: PPARα/PPARγ dual agonist, PAR5359 ameliorates DSS-induced colitis in mice.** (**A**) Schematic summarizing the workflow for testing PPARα/γ dual agonist (PAR5359) in DSS-induced colitis. Briefly, mice were fed with 2.5% DSS in drinking water for 7 days followed by 2 days of normal drinking water. PAR5359 was given through intrarectal route using an oral gavage needle. The tip of oral gavage needle was greased with medical grade ointment for easy and safe administration. On 9$^{th}$ day all group mice were sacrificed, and colons were analyzed for its morphology, histology and gene expression (RNAseq and qPCR). (**B**) Line graphs display daily weight of mice, from the day of DSS administration (day 1) to the day of sacrifice (day 9). (**C**) Line graphs display disease activity index (DAI) scores, calculated for the days 5, 7 and 9 after DSS administration, which accounts for stool consistency (0-4), rectal bleeding (0-4), and weight loss (0-4). (**D**) Scatter plots with bar graphs display the length of the excised colon at sacrifice (day 9). (**E**) Images representative of H&E-stained sections of the distal colon are shown (**F**) Bar graphs showing histological score of H&E sections mentioned in E. White arrowheads = immune cell infiltrates. Statistics: All results are displayed as mean ± SEM. Significance was tested using two-way/one-way ANOVA followed by Tukey's test for multiple comparisons. Significance: *, $p < 0.05$; **, $p < 0.01$, ***, $p < 0.001$, ****, $p < 0.0001$.

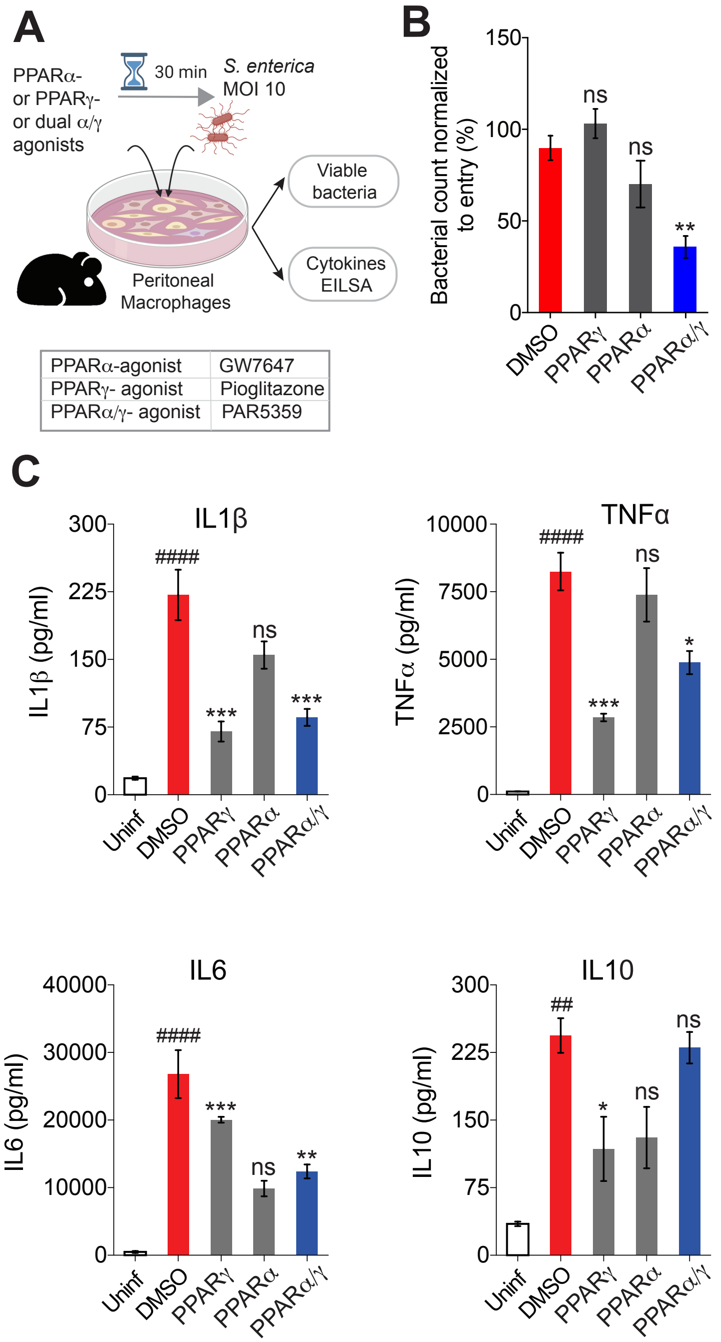


**Supplementary Figure 10: PPARα and PPARα/γ-dual agonists enhance, whereas PPARγ agonist delay bacterial (*Salmonella enteritica*) clearance.** (**A**) Schematic displays the experimental design and workflow. Thioglycolate-induced murine peritoneal macrophages (TG-PM) pretreated with PPAR agonists (see box, below; 20 nM GW7647, 10 μM Pioglitazone and 1 μM PAR5359) were infected with *Salmonella enterica* (*MOI* 10) and subsequently analyzed at 6 h post-infection for bacterial count (Gentamicin protection assay) and secretion of inflammatory cytokines (in supernatant media by ELISA). (**B**) Bar graphs show percent internalized viable bacterial counts of *Salmonella enterica.* (**C**) Bar graphs display the extent of secreted cytokines (IL1β, IL6, TNFα and IL10) in the supernatant media. Statistics: One-way ANOVA followed by Tukey's test for multiple comparisons was performed to test significance. '#' significance over uninfected TG-PMs and '*' shows significance over *Salmonella*-infected cells. All results are from at least three independent experiments and results displayed as means ± SEM. ns, non-significant, *, $p < 0.05$; **, $p < 0.01$, ***, $p < 0.001$.

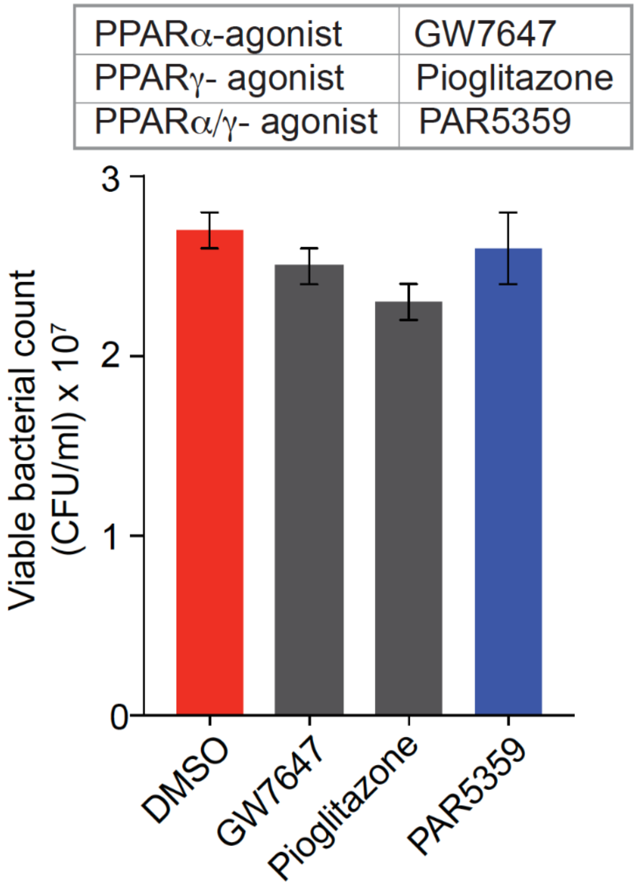


**Supplementary Figure 11: PPAR agonists did not affect the viability of AIEC- LF82.** *AIEC*-LF82 bacteria (~2 x $10^7$) were incubated for 1 h with RPMI media containing 10% FBS and PPAR agonists (see box, below; 20 nM GW7647, 10 mM Pioglitazone and 1 mM PAR5359) at 37°C in CO2 incubator. The culture media was subsequently analyzed for the bacterial count. All results are from three independent experiments and results are displayed as means ± SEM. Significance was tested using one-way ANOVA.

#### TABLES

**Table S1: Table summarizing studies to date and their claims regarding the protective role of PPARγ in IBD**

| **Interventions/Study models** | **Outcome / Major conclusions** | **Reference** |
|---|---|---|
| DSS-induced colitis model in WT and Intestinal endothelial cells (IEC; Villin Cre)-specific PPARγ KO mice<br>Agonist: Rosiglitazone | DSS-induced colitis is worsened in KO mice, compared to WT littermates. PPARγ expressed in the IEC has an endogenous role in protecting against colitis. | (29) |
| DSS-induced colitis model in WT and CD4+ T cell-specific PPARγ KO mice | DSS-induced colitis is worsened in KO mice, compared to WT littermates. PPARγ in T cells is involved in preventing gut inflammation by regulating adhesion molecules and inflammatory mediators. | (30) |
| DSS-induced colitis model in WT and Macrophage-specific PPARγ KO mice<br>Agonist: Pioglitazone | Macrophage-specific PPARγ KO exacerbated DSS-induced colitis, impaired Treg compartment, and increased LP CD8+ T cells. In addition, the protective effect of Pioglitazone, the presence of PPARg in macrophages is required. | (31) |

**Table S2: Table summarizing studies to date and their conflicting claims regarding the role of PPARα in IBD**

| Sl. No. | Interventions/Study models | Outcome / Major conclusions | Reference |
|---|---|---|---|
| 1. | Dinitrobenzene sulfonic acid (DNBS) induced IBD in WT and PPARα$^{-/-}$ mice.<br>With or without PPARα agonists WY-14643. | DNBS-induced colitis is worsened in KO mice, compared to WT littermates.<br>PPARa and its agonist WY-14643 protects from IBD.<br>**PPARα is protective** | (32) |
| 2. | DSS-induced colitis in Interleukin-10 knockout (IL-10$^{-/-}$)<br>Agonist: Fenofibrate | **PPARα is protective** | (33) |
| 3. | DNBS-induced IBD in WT and PPARα$^{-/-}$ mice, and treatment with dexamethasone, a synthetic glucocorticoid | PPARα enhances the anti-inflammatory effect of dexamethasone.<br>**PPARα is protective** | (34) |
| 4. | DSS-induced colitis in mice treated with<br>Agonist: WY14643 | **PPARα is protective** | (35) |
| 6. | DSS-induced colitis in PPARα$^{-/-}$ and WT mice<br>Agonist: WY14643, | PPARα agonist worsens colitis in a PPARα-dependent manner<br>**PPARα is protective** | (36) |
| 7. | DSS and TNBS-induced colitis in WT and PPARα$^{-/-}$mice and *Salmonella typhi* induced colitis, with or without treatment with PPARα agonist.<br>Agonist: Fenofibrate | Increased inflammation in WT, but not KO mice.<br>PPARα agonist worsens colitis in a PPARα-dependent manner<br>**PPARα is harmful** | (37) |
| 9. | DSS-induced colitis in WT and PPARα$^{-/-}$mice; treatment with PPAR agonist<br>Agonist: fenofibrate | Colitis worsened by agonists in WT, but not KO mice.<br>PPARα agonist worsens colitis in a PPARα-dependent manner<br>**PPARα is harmful** | (38) |

**Table S3: PPARα/γ dual agonists, their potency and market status.**

| PPARα/γ Dual Agonist | EC50 (α) | EC50 (γ) | Status |
|---|---|---|---|
| Muraglitazar | 320.0nM<br>5680nM | 110.0nM<br>243nM | Discontinued (39, 40) |
| Tesaglitazar | 4780nM<br>1200nM | 3420nM<br>1300nM | Discontinued (40, 41) |
| Naveglitazar | 2816nM | 361nM | Discontinued (42) |
| Ragaglitazar | 3200nM | 600nM | Discontinued (43) |
| Farglitazar | 250nM<br>450nM | 0.2nM<br>0.34nM | Discontinued (44, 45) |
| Imiglitazar | 8nM | 4nM | Discontinued (46) |
| Netoglitazone | 100nM[9] | 3000nM[9] | Discontinued (47) |
| Reglitazar | 1900nM | 83nM | Discontinued (48) |
| MK0767 | 140nM | 83nM | Discontinued (49) |
| KRP-297 | 850nM | 83nM | Discontinued (50) |
| TZD18 | 26nM | 14nM | Preclinical (51) |
| Chiglitazar | 1200nM | 80nM | Phase II clinical trials (52) |
| Aleglitazar | 50nM<br>5nM | 21nM<br>9nM | Phase III clinical trials (40, 53) |
| PLX429 | - | - | Preclinical |
| AVE0847 | - | - | Phase II clinical trials |
| Azaindole-α-alkyloxyphenylpropionic acid | - | - | Preclinical |
| BVT-142 | - | - | Preclinical |
| O-Arylmandelic acid derivatives | - | - | Preclinical |
| Amide substituted with α-substituted-β-phenylpropionic acid derivatives | - | - | Preclinical |
| 2-Alkoxydihydro cinnamate derivatives | - | - | Preclinical |
| LY51029 | - | - | Preclinical |
| α-Aryloxyphenyl acetic acid derivatives | - | - | Preclinical |
| Tricyclic-α-alkyloxyphenyl propionic acids | - | - | Preclinical |
| Saroglitazar | 0.00065nM | 3nM | Phase II clinical trials,<br>Approved in India and Mexico (54) |

**Table S4: PPAR agonists and antagonists used in this study.**

| Common name | Chemical name | structure | $EC_{50}$Potency | Tested in IBD | Reference |
|---|---|---|---|---|---|
| **GW7647** PPARα agonist | 2-4-2-4 Cyclohexylbutyl)(cyclohexylcarbamoyl) amino]ethyl}phenyl sulfanyl]-2-methylpropanoic acid | 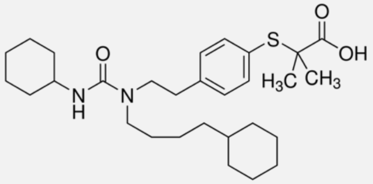 | PPARα agonist: 2.1± 0.05 nM | No | (55) (56) |
| **Pioglitazone** PPARγ agonist | 5-[4-[2-[5-ethylpyridin-2-yl]ethoxy]benzyl]thiazolidine-2,4-dione | 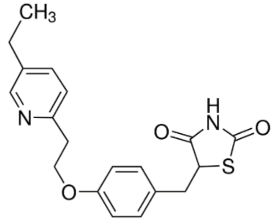 | PPARγ agonist: 1.1± 0.03 mM | Yes | (57) |
| **PAR5359** PPARα/γ dual agonist | 3-(4-{2-[4-(4-Chloro-phenyl)-3,6-dihydro-2H-pyridin-1-yl]-ethoxy}-phenyl)-2-ethoxypropionic acid | 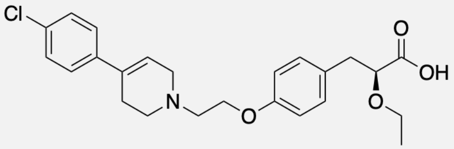 | PPARα agonist: 0.16± 0.005 µM PPARγ agonist: 0.12± 0.006 µM | No | (58) |
| **GW 6471** PPARα antagonist | *N*-((2*S*)-2-(((1*Z*)-1-Methyl-3-oxo-3-(4-(trifluoromethyl)phenyl)prop-1-enyl)amino)-3-(4-(2-(5-methyl-2-phenyl-1,3-oxazol-4-yl)ethoxy)phenyl)propyl)propanamide | 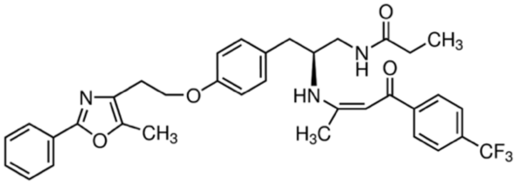 | PPARα antagonist: 0.24 mM | No | (59) (60) |
| **GW 9662** PPARγ antagonist | 2-Chloro-5-nitro-N-phenylbenzamide | 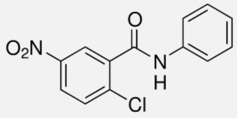 | PPARγ antagonist: 3.3 nM | No | (61) |

**Table S5: Characteristics of IBD patients used as source of PBMCs for use in bacterial clearance assays.**

| Patient code | Sub-disease | Gender | Treatment history |
|---|---|---|---|
| **CD 76** | Inflammatory | Male | Infliximab current; No prior Biologics |
| **CD77** | Stricturing | Male | Infliximab in the past |
| **CD79** | Penetrating | Male | Infliximab in the past |
| **CD80** | Stricturing | Female | Adalimumab current; No prior Biologics |
| **CD81** | Penetrating | Female | Infliximab current; Previoulys on Cimzia and Stelara |
| **CD 82** | Inflammatory | Female | Biologic Naïve |
| **CD 83** | Stricturing, Penetrating | Male | Infliximab current; No prior Biologics |
| **UC72** | UC | Female | Biologic Naïve |
| **UC73** | UC | Female | Biologic Naïve |
| **UC 74** | UC | Female | Biologic Naïve |
| **UC 75** | UC | Female | Biologic Naïve |
| **UC 76** | UC | Female | Off therapy, previous had one dose of Entyvio |
| **UC 77** | UC | Female | Adalimumab; No prior Biologics |
